# *In Situ* Landscape of Focal Adhesions and Cytoskeletal Integration Revealed by Cryo-Electron Tomography

**DOI:** 10.64898/2026.04.08.717201

**Authors:** Piao Yu, Lingyun Zhao, Ashraf Al-Amoudi, Stefan T. Arold

## Abstract

Focal adhesions (FAs) are dynamic hubs for mechanotransduction, linking the extracellular matrix to actin fibers, intermediate filaments, and microtubules. Using cryo–electron tomography combined with 3-dimensional segmentation and subtomogram averaging, we visualize *in situ* the structural architecture of the FA environment at the leading edge of human fibroblasts. Our analysis reveals a rich architectural diversity within the FA landscape, where the spatial organisation and interplay of FA protein clusters, actin, vimentin, and microtubules change from the actin bundle core to its tip and across adjacent regions. Notably, we reveal diverse arrangements and connections of vimentin filaments, supporting their multifaceted role in the control and mechanics of adhesions. Together, these findings establish a structural framework for FA maturation and cytoskeletal integration, extending classical lamellipodial adhesion models and providing mechanistic insight into how FAs coordinate force transmission during cell migration.

## INTRODUCTION

Focal adhesions (FAs) generate traction forces that enable cell migration by dynamically assembling and disassembling during movement. Beyond their mechanical role, FAs act as signaling hubs that allow cells to sense and respond to extracellular mechanical cues^1^. These dynamic structures, typically 0.25–20 µm in length and ∼0.1 µm in height, physically link the actin cytoskeleton to the extracellular matrix (ECM). Integrins, specifically the αVβ3 heterodimer, initiate FA assembly by coupling the ECM to intracellular scaffolds and signaling networkss^2,3^. Mature FAs comprise numerous copies of hundreds of proteins, with paxillin, focal adhesion kinase (FAK), kindlin, vinculin, and talin among their core components^4,5,6^. Beyond anchoring the actin cytoskeleton to the extracellular matrix, FAs also process and transduce signals, including those controlling autophagy and cell death after detachment (anoikis)^1,7^. Their dynamic nature is essential for embryogenesis, wound healing, and synaptic plasticity. Accordingly, FA dysregulation is linked to cancer metastasis and neurodegenerative disorders^8,9,10^.

During FA maturation, interactions between talin and KN motif and ankyrin repeat domain-containing (KANK) proteins create a mechanically sensitive linkage between the integrin–talin complex and the cortical-microtubule-stabilizing complex^11,12^. As actomyosin contractility increases the tension transmitted through the integrin–talin–actin axis, talin undergoes force-induced unfolding^4^, which disrupts its interaction with KANK^11,12^. This mechanical separation generates distinct tension domains: a high-tension FA core that excludes KANK, and a peripheral low-tension FA belt enriched in KANK-dependent cortical–microtubule–stabilizing complexes. This low-tension zone maintains microtubule coupling and locally facilitates adhesion disassembly by providing compressive resistance^11,12,13,14^.

Alongside actin filaments and microtubules, intermediate filaments constitute the third major cytoskeletal system involved in FAs. Intermediate filaments are ∼10 nm in diameter and built from α-helical coiled-coil dimers that assemble into staggered, antiparallel tetramers^15^. In contrast to the polarized and dynamic actin and microtubule networks, intermediate filaments are non-polar, highly deformable, and optimized for mechanical resilience rather than motility^16^. Their molecular composition is cell-type specific, where keratins are dominant in epithelial cells, desmin in muscle, and vimentin in mesenchymal cells^16^. Intermediate filament–FA interactions exert bidirectional control over adhesion dynamics. Anchoring of vimentin to mature FAs or fibrillar adhesions stabilizes high-tension adhesions and suppresses turnover^17^, whereas loss of intermediate filament anchorage disrupts mechanotransduction and produces dysfunctional, low-tension adhesions that also turn over inefficiently due to impaired disassembly^18,19^. Beyond their direct FA engagement, intermediate filaments integrate into both the actomyosin and microtubule networks, reinforcing cytoskeletal connectivity and buffering mechanical stress at adhesion sites. Through combined participation in actomyosin-driven tension and microtubule-supported compression, intermediate filaments enhance cellular viscoelastic resilience and modulate local stress responses^20^. Together, the actin, microtubule, and intermediate filament form a mechanically coupled, adaptive cytoskeletal continuum that orchestrates FA mechanosensitivity, turnover, and renewal.

The spatial arrangement of FA sites has been investigated by different approaches. Super-resolution analyses have revealed that the FA complexes under the actin bundle core are vertically stratified across ∼40 nm, comprising signaling, force-transducing, and cytoskeletal layers^6,12,21^. Atomic force microscopy–based three-dimensional reconstructions of FA topography further revealed a wedge-shaped architecture, in which finger-like microfilaments at the leading edge project about 100 nm above the substratum in a shallow ∼3° angle^21^. Additionally, cryogenic electron tomography (cryo-ET) was used to structurally characterize specific actin-associated particles^22,23^. However, the three-dimensional integration of FAs with surrounding cytoskeletal systems (i.e., actin, intermediate filaments, and microtubules) has not yet been resolved *in situ*.

## RESULTS

### FA-peripheral densities next to the actin bundle interface with actin filament and intermediate filament networks

To localize FA sites for cryo-ET, we established a semi-correlative fluorescence and electron microscopic workflow (**Supplementary Fig. 1**, route 1). Fibroblasts were grown on gold 200-mesh holey carbon finder grids supplemented with an additional 2 nm carbon layer to enable cell adhesion, followed by chemical fixation and immunolabeling with an Alexa Fluor 488-conjugated anti-vinculin antibody. The grids were imaged by fluorescence microscopy and subsequently vitrified for cryo-ET analysis. Tomogram acquisition sites were selected by matching fluorescence footprints at the cell leading edge to the corresponding grid coordinates (**Fig. 1**, **Table 1**).

**Figure 1.**
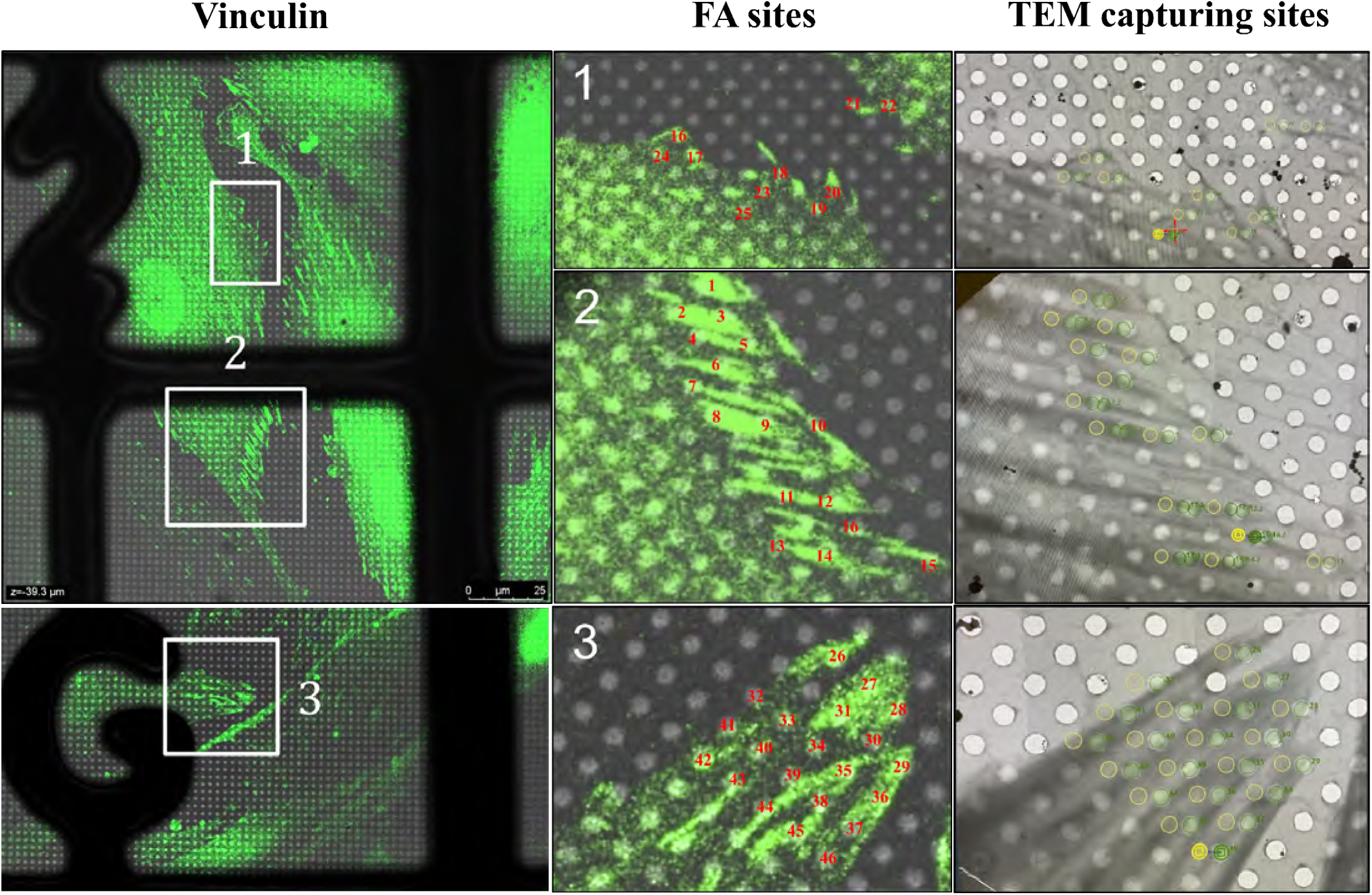
Correlative identification of vinculin-positive FA sites for cryo-ET data collection. Fluorescence images of fibroblasts immunolabeled with Alexa Fluor 488-conjugated anti-vinculin antibody (left) show the overall FA distribution across EM grids. Boxes (1–3) indicate selected FA-enriched areas. The middle panels show magnified views of these boxes, with individual FA sites (numbered in red) chosen for tomography acquisition. The right panels display corresponding low-magnification TEM images used to locate the same FA sites on the grid. Green circles indicate the positions selected for tilt-series collection.

A consistent observation in the fluorescence images was that vinculin-positive FA plaques were predominantly located at the cell leading edge, where prominent actin bundles are known to form (**Fig. 1**). Based on the relative position of FA plaques to the actin bundle, we defined two regions for tomogram acquisition: The tip region, which corresponds to FA plaques at the tips of actin bundles at the cell leading edge, and the core region, corresponding to the FA plaques posterior to the tip region (**Fig. 2a, Supplementary Fig. 2**). Two-dimensional projection micrographs of the putative actin bundles matched the fluorescence signals (**Supplementary Fig. 2b,** white and red dashed outline), confirming that the fluorescent plaques were positioned on the bundled actin area rather than in the adjacent cytoplasm.

**Figure 2.**
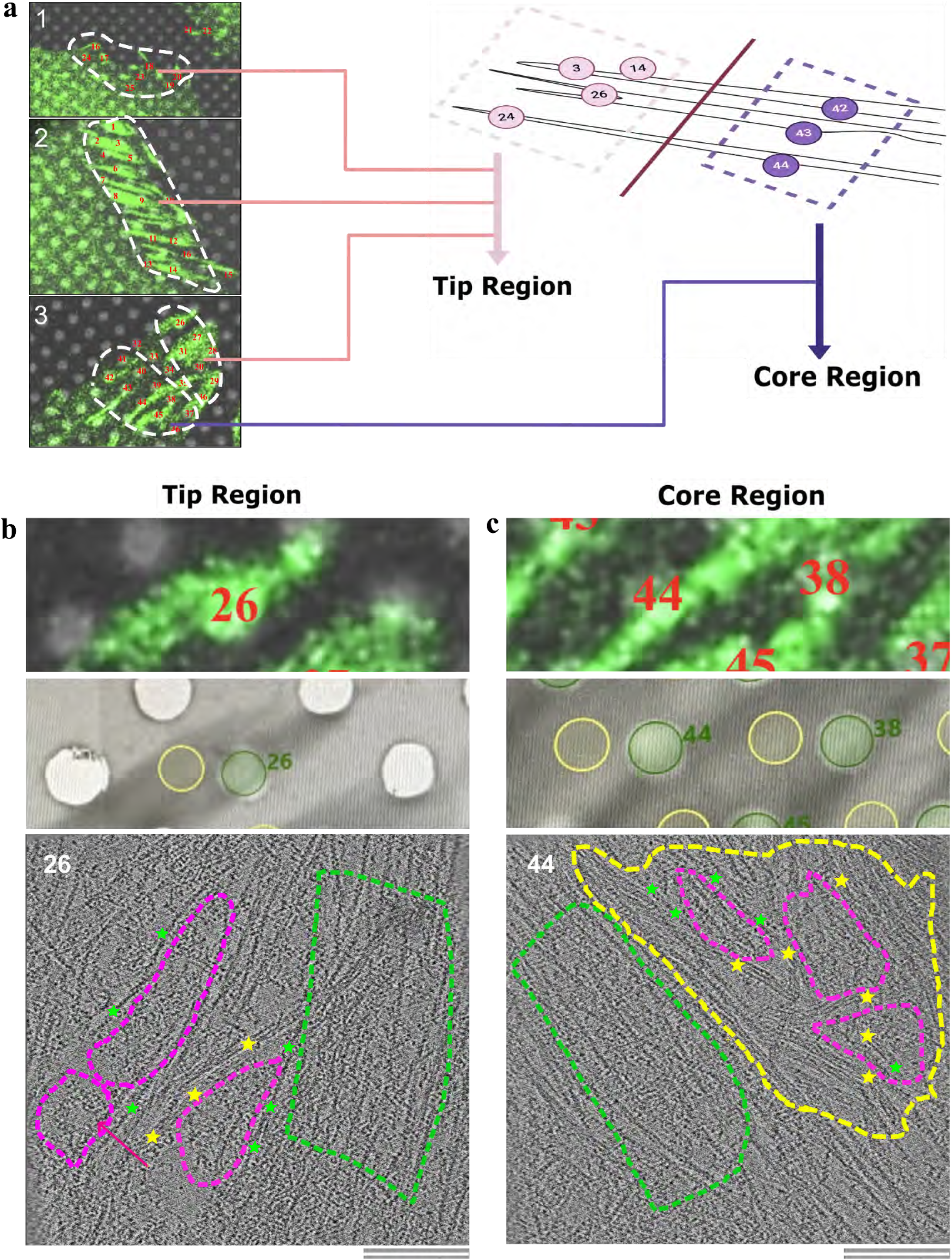
Correlative fluorescence–cryo-ET analysis of FA sites at Tip Region and Core Region. (a) Left, representative fluorescence image of FA sites on the cryo-EM grid (same FA-site column as shown in Fig. 1). Boxes (1–3) indicate selected FA-enriched areas. White dashed outlines denote FA-enriched regions, which were classified as tip regions or core regions. Right, schematic illustrating the relative spatial relationship between the FA-enriched regions and the filament bundle. Pink- and purple-shaded circles represent sampling positions assigned to the tip region and core region, respectively. Numbers within the circles correspond to the position indices of the sampling locations shown in the FA-enriched areas on the left. Black lines outline the overall shape of the thick filament bundle at the cell leading edge. (b,c) Correlative fluorescence–cryo-ET views from representative sampling positions in the tip region (position 26; b, left column) and the core region (position 44; c, right column). For each column, the top panel shows fluorescence localization of FA sites, the middle panel shows the corresponding cryo-EM grid view with circles marking the selected positions, and the bottom panel shows a representative tomographic slice from the indicated sampling position. The magenta arrow highlights a protein density associated with an actin filament. Magenta dashed outlines indicate other dispersed protein densities. Green dashed outlines denote dense actin-bundle regions, whereas yellow dashed outlines mark intermediate filament–like rich regions. Green stars and yellow stars indicate actin filaments and intermediate filament–like filaments located near the protein densities, respectively. Scale bar, 100 nm.

Cryo-ET datasets acquired from different sites within the tip region showed distinct ultrastructural features. The body of the actin bundle consisted of densely packed and well-aligned actin filaments (**Fig. 2b, green dashed outline**). The periphery of the actin bundle exhibited more disordered and sparse actin architectures (**Supplementary Fig. 2c,d, green star**). The lateral side of the actin bundle harboured sparse, discontinuous protein densities connected to the bundle through individual actin filaments (**Fig. 2b, magenta dashed outline**). This flanking area also occasionally contained additional fibers, distinct from actin filaments and microtubules (**Fig. 2b, yellow star**). Based on their intermediate size and apparent flexibility, we tentatively interpreted these structures as intermediate filaments.

Cryo-ET of the actin bundle core region showed intermediate filament–rich structures in the area flanking the dense actin bundle (**Fig. 2c, Supplementary Fig. 2e, f; yellow dashed outline**). At the interface between actin and intermediate filament (**Supplementary Fig. 2f**), or within the intermediate filament–rich region itself (**Fig. 2c**), we observed sparse and discontinuous protein densities connected to individual actin filaments (**Fig. 2c, Supplementary Fig. 2f; magenta dashed outline)**, akin to the tip region.

We concluded that fluorescence-guided targeting enabled the localization of vinculin-positive FA zones and direct visualization of their local cytoskeletal architecture. Along the lateral periphery of the actin bundles, we observed sparse and discontinuous protein densities in direct contact with individual F-actin filaments and neighboring intermediate filaments, indicating these densities engage adhesion structures through interactions with both actin and intermediate-filament cytoskeletal elements. Based on their location at the bundle periphery, we herein refer to these densities as FA-peripheral densities. Notably, and in contrast to the canonical view of FA organization^6^, we did not detect large and continuous FA densities directly under the actin bundle (termed herein FA-core densities) in these tomograms of intact cells.

### Immunogold labeling identifies FA-peripheral densities as vinculin-containing FA assemblies

To enhance image contrast and reduce sample crowding, we next developed a protocol to remove the plasma membrane and non-adherent components while preserving the architecture of the FA sites (**Supplementary Fig. 1, route 2)**. This procedure was adapted from previously described detergent-based membrane extraction protocols^23,24^ to reinforce cellular adhesion structures in our samples.

Cryo-ET visualization of these samples confirmed that they retained the hallmarks of FAs, including actin bundles, microtubules and FA-peripheral protein clusters (**Supplementary Fig. 3, Table 1**). However, the treatment increased the image contrast, as expected from the thinner membrane-less sample. Tomograms collected from the actin-bundle body of the tip region revealed parallel actin filaments. Some filaments were connected by ∼30-nm linear densities (**Supplementary Fig. 3a, Magenta arrow**), consistent with α-actinin crosslinkers^12,25^. Tomograms collected from the flanks of the actin bundle tip additionally showed microtubules (**Supplementary Fig. 3b, Magenta star**) and FA-peripheral densities directly contacting individual F-actin filaments adjacent to the actin bundle (**Supplementary Fig. 3b, yellow star**).

We next applied immunogold labeling against vinculin to the membrane-less samples as a spatial marker for FA localization (**Supplementary Fig. 1**, **route 3**). We acquired tomograms of gold-positive positions at either the actin bundle tip, core, or periphery (**Fig. 3**). The ultrastructural features of the gold-labeled sites, specifically the spatial arrangement of actin filaments, intermediate filaments, microtubules, and dispersed FA-peripheral densities contacted by individual actin filaments (**Fig. 4; Supplementary Fig. 4**), were consistent with those observed in whole-cell or adhesion-focused tomograms, confirming that vinculin-associated gold particles reliably mark FA sites.

**Figure 3.**
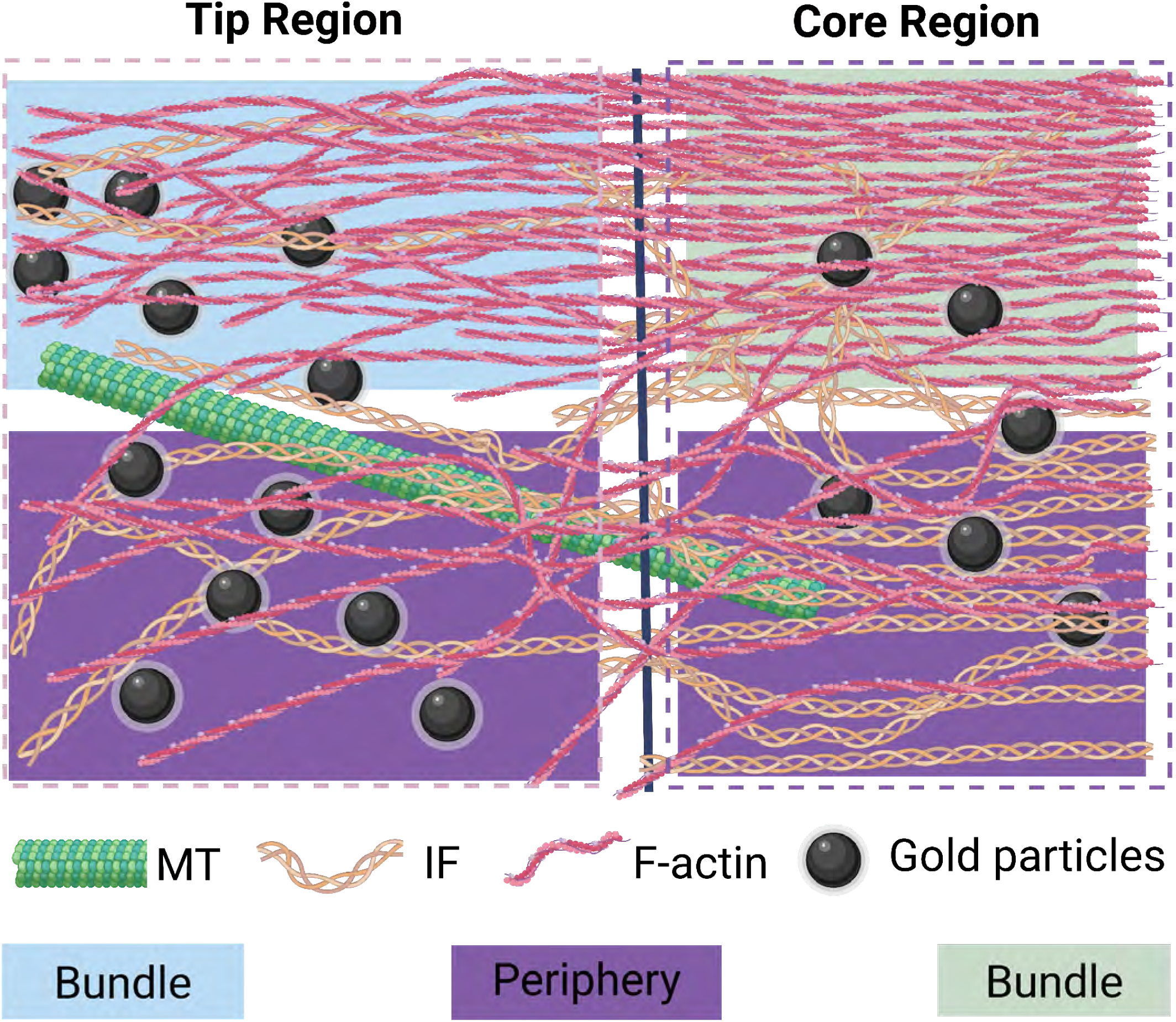
Sampling positions for cryo-ET at actin-bundle and periphery regions. Schematic showing tip and core sampling sites with their characteristic cytoskeletal features. Background shading delineates the bundle zones (blue for tip, green for core) and their periphery regions (purple), which guided site selection during cryo-ET acquisition. Gold particles (black spheres) mark immunogold-defined FA sites. Microtubules (MTs, green cylinders), intermediate filaments (IFs, beige helices), and actin filaments (pink filaments) are shown together with gold fiducial particles (black spheres).

**Figure 4.**
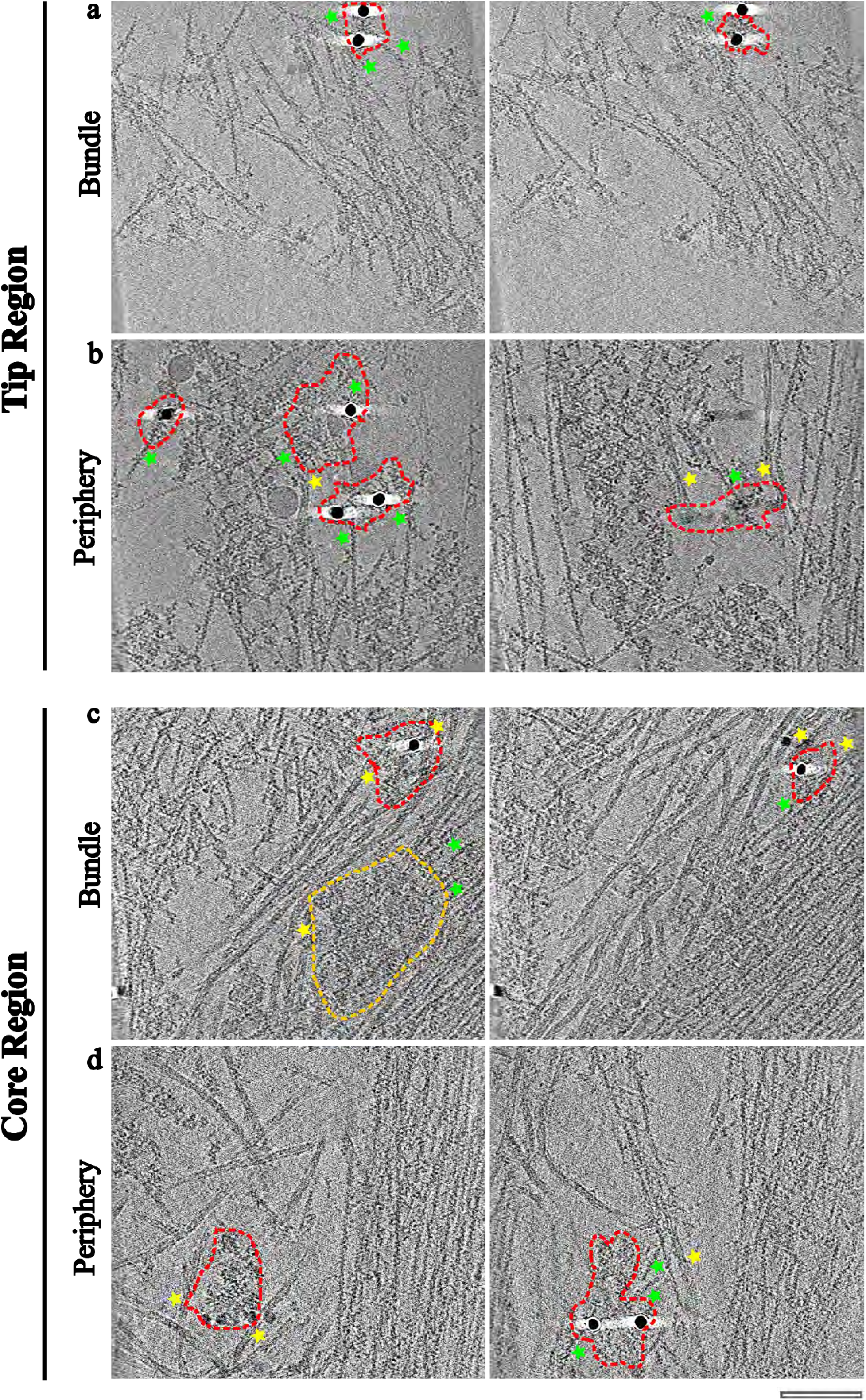
Representative tomographic slices showing FA densities at bundle and periphery regions. Representative tomographic slices were extracted from sampling positions defined in the schematic shown in Fig. 3. Each row shows two representative slices from the same tomogram, highlighting the structural relationships between protein densities and surrounding cytoskeletal filaments within the selected region. Panels (a,b) show sampling positions from the tip region, whereas panels (c,d) show positions from the core region. Within each region, bundle areas are shown in panels (a) and (c), and periphery areas are shown in panels (b) and (d). Gold particles mark FA locations, at which either FA-peripheral protein densities (red dashed outlines) or classical FA-core densities (yellow dashed outlines) are observed. MTs (red stars), IFs (yellow stars), and actin filaments (green stars) are observed either contacting or in proximity to these gold-labeled densities. More comprehensive series of tomographic slices spanning low to high Z positions for the same sampling sites are shown in Supplementary Fig. 4. Scale bar, 100 nm.

At the actin bundle tip region, cryo-ET revealed gold particles colocalized with dispersed FA-peripheral densities within the thinning actin bundle (**Fig. 4a**, red **dashed outline**). These gold particles were positioned adjacent to single actin filaments (**Fig. 4a, green star**) that extended obliquely into FA-peripheral densities. Neighboring actin filaments were accompanied by intermediate filaments running alongside them (**Supplementary Fig. 4a, panel 2–4; Supplementary Fig. 4b, panel 5; green and yellow stars**). One microtubule was visible at the periphery of the actin bundle (**Supplementary Fig. 4a, panel 1; red star**).

Tomograms taken on the lateral periphery of the tip region revealed a cytoskeletal organization similar to that of the tip central region, but with more dispersed FA-peripheral densities (**Fig. 4b; Supplementary Fig. 4c-d**). Vinculin-labeled gold particles also localized to these dispersed densities (**Fig. 4b, Supplementary Fig. 4c-d; red dashed outline**), which were contacted by both actin filaments and intermediate filaments (**Fig. 4b, Supplementary Fig. 4c-d; green and yellow stars**).

Cryo-ET of the actin bundle core regions revealed gold particles localized at the base of the dense actin bundle, where large, continuous FA-core densities were present (**Fig. 4c**, **Supplementary Fig. 4e; yellow dashed outline**). Hence, the membrane-less tomograms showed the canonical large FA-core densities beneath the dense actin bundle which were not detectable in tomograms of intact cells. Tomograms of the periphery of the actin core region further showed gold labeling of dispersed FA-peripheral densities positioned at the interface of the actin–intermediate filament fibers (**Fig. 4c**, **Supplementary Fig. 4e; red dashed outline**), and within the intermediate filament region itself (**Fig. 4c-d**, **Supplementary Fig. 4e-f; red dashed outline**). Both FA-peripheral and FA-core densities were observed in direct contact with actin and intermediate filaments (**Fig. 4c-d**, **Supplementary Fig. 4e-f; green and yellow stars**). These results indicated that the FA assembly beneath the dense actin bundle occurs at the intersection of both cytoskeletal systems of F-actin and intermediate filaments.

In conclusion, gold labeling marked both the FA-core densities and the laterally distributed FA-peripheral densities, confirming both as vinculin-containing FA types despite their distinct spatial distribution. Moreover, the FA-peripheral densities were frequently connected to FA-core densities through F-actin filaments that extended from the actin bundle into the FA-core region.

### In Situ Structural Support for the Intermediate Filament being Vimentin

Although fibroblasts predominantly express vimentin^16^, we sought to confirm that the intermediate filaments we observed at FA sites corresponded to vimentin. To this end, we performed subtomogram averaging on 28 tomograms enriched in these filaments (**Supplementary Fig. 5, Table 1**).

The intermediate filaments were first traced in Dynamo^26^ and 37,582 subtomograms were extracted with a 6 nm overlap and a 38 nm box size. To obtain higher-resolution insight into the 3D structure of the intermediate filaments, these subtomograms were then projected at 13 nm thickness using the Actin Polarity Toolbox (APT)^27^ and the resulting 2D projections were subjected to several rounds of unsupervised 2D classification in RELION (**Supplementary Fig. 6**) ^15,28^. Good 2D class averages, combined with initial priors for the in-plane rotation and tilt angle saved in the extracted subtomograms from Dynamo, were then exported to cryoSPARC for 3D refinement. The resulting 3D reconstruction yielded maps at a resolution of 16 Å using homogeneous refinement (**Supplementary Fig. 7a**). Helical refinement, for which we determined the helical rise (40 Å) and twist (72°), improved the resolution to 13 Å (**Supplementary Fig. 7b,c**). The helical rise and twist were in agreement with those determined previously for vimentin^15^. Both maps revealed a tubular architecture ∼11 nm in diameter with a central lumen. Cross-sections were consistent with the five-protofibril organization characteristic of vimentin filaments (**Fig. 5a,b**). Rigid-body fitting of the vimentin filament model (PDB 8RVE) into the 12.96 Å map produced a cross correlation coefficient of 0.86. The atomic model followed filament densities in the side view **(Fig. 5c**), and its five-protofibril arrangement matched the five wall densities in cross-section (**Fig. 5d-e**). The lumen density observed in the reconstruction was also compatible with the amyloid-like fiber formed by vimentin low-complexity domains between protofibrils^15^. Taken together, the *in situ* filament morphology, helical rise and twist, GSFSC-validated reconstructions, and structural fit collectively support that these intermediate filaments are mature vimentin fibers.

**Figure 5.**
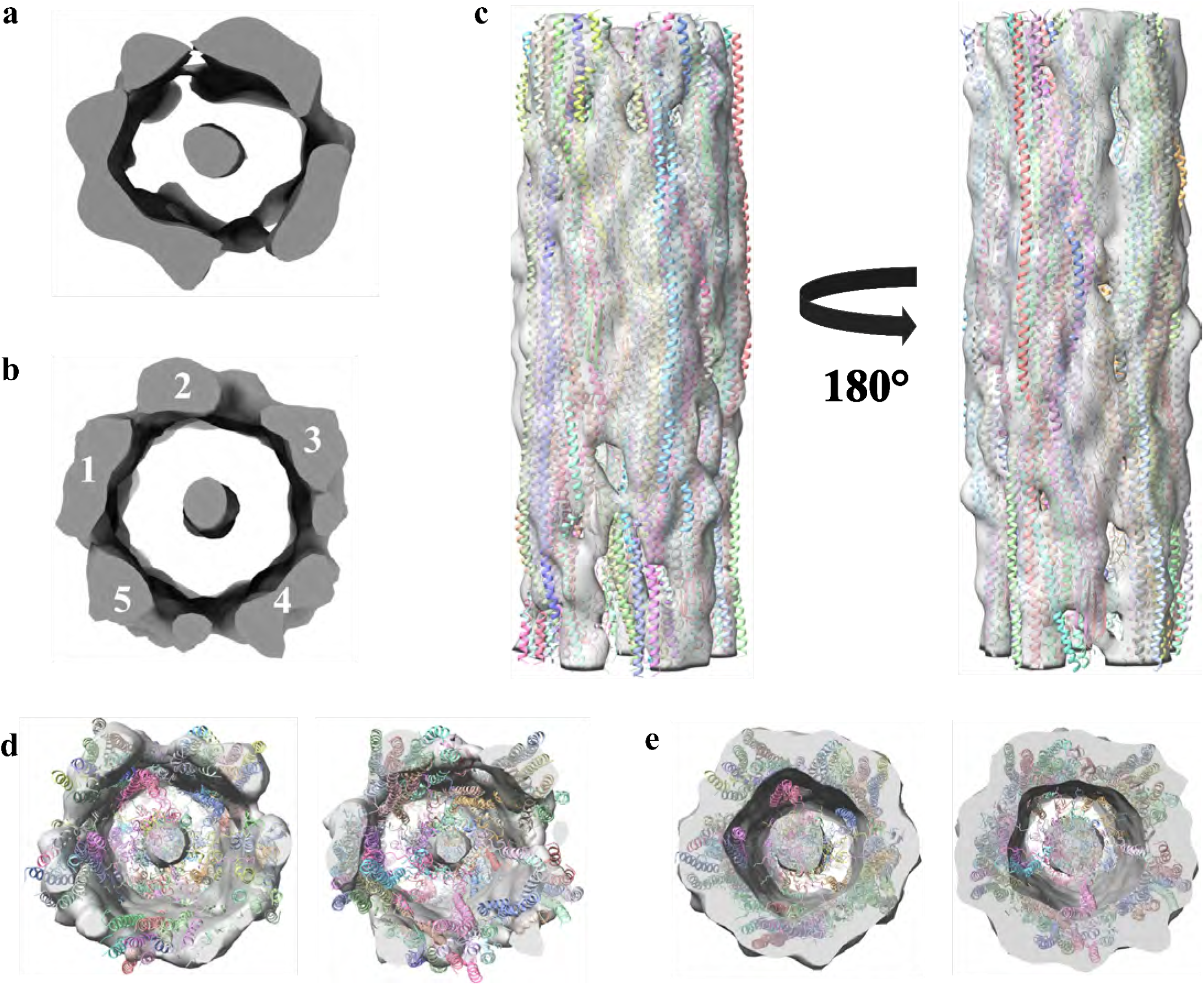
Structural identification and characterization of IFs. (a,b) CryoSPARC 3D reconstructions of vimentin intermediate filaments obtained by homogeneous refinement (a) and helical refinement (b), respectively. The helical refinement resolves continuous longitudinal density features along the filament axis that are not apparent in the homogeneous reconstruction. (c) Side views of the helical refinement map fitted with atomic models of vimentin (PDB 8RVE), showing the alignment of five longitudinal density ridges along the filament axis. The two views are related by a 180° rotation around the Z axis. The atomic model was fitted as a rigid body into our EM density map using ChimeraX. The fit achieved a cross-correlation coefficient (CCC) of 0.86, indicating a strong agreement between the model and the map. (d,e) End-on views of the fitted model displayed at two different map contour levels. Five protofibrils correspond to the densities forming the outer wall of the filament. An additional central density within the lumen is observed, consistent with amyloid-like fibers formed by the low-complexity domains of vimentin, as previously reported^15^.

### Stratified spatial organization of FA-peripheral densities and cytoskeletal networks at the periphery of the core region

Immunogold-labeled samples confirmed that vimentin fibers flanking the dense actin bundles contained dispersed FA-peripheral assemblies (**Fig. 4c,d**). To resolve the three-dimensional spatial relationships between these assemblies and the surrounding cytoskeletal networks, we acquired cryo-ET datasets from the vimentin-rich periphery of the actin bundle core. During data collection, the dense actin core bundles were intentionally excluded from the field of view. We first acquired low-magnification search maps to determine the bundle orientation and identify the tip/periphery boundaries, enabling precise positioning of each tilt series. This ensured that the resulting tomograms captured sufficient peripheral volume to visualize FA-peripheral densities within their local cytoskeletal context. Because our earlier analyses had established that these densities localize to the bundle periphery, re-imaging the dense core at high resolution was unnecessary.

A representative tomogram from these acquisitions (**Fig. 6**) shows vinculin-associated gold particles positioned lateral to the dense actin bundle, embedded within a surrounding network of vimentin filaments. Consecutive tomographic Z-slices spanning from membrane-proximal region upward into the cytoplasm revealed a gradual reorganization of cytoskeletal elements. In the membrane-proximal region, vimentin filaments formed a regularly aligned array running nearly parallel to the dense actin bundle **(Fig. 6**, 1; yellow dashed line). With increasing distance from the membrane, this ordered array progressively transitioned into a heterogeneous meshwork of intersecting vimentin filaments and the FA-peripheral proteins resided in the spaces between them (**Fig. 6**, 5–8). At intermediate height, a microtubule became visible, oriented parallel to the actin bundle (**Fig. 6**, 5–7). A sparse layer of F-actin appeared above the FA-peripheral protein density in direct contact with this region (**Fig. 6**, 8). We measured the diameters of the microtubule and vimentin filaments to be approximately 24 nm and 11 nm, respectively (**Supplementary Fig. 8b**), consistent with their reported morphologies^15,29,30^.

**Figure 6.**
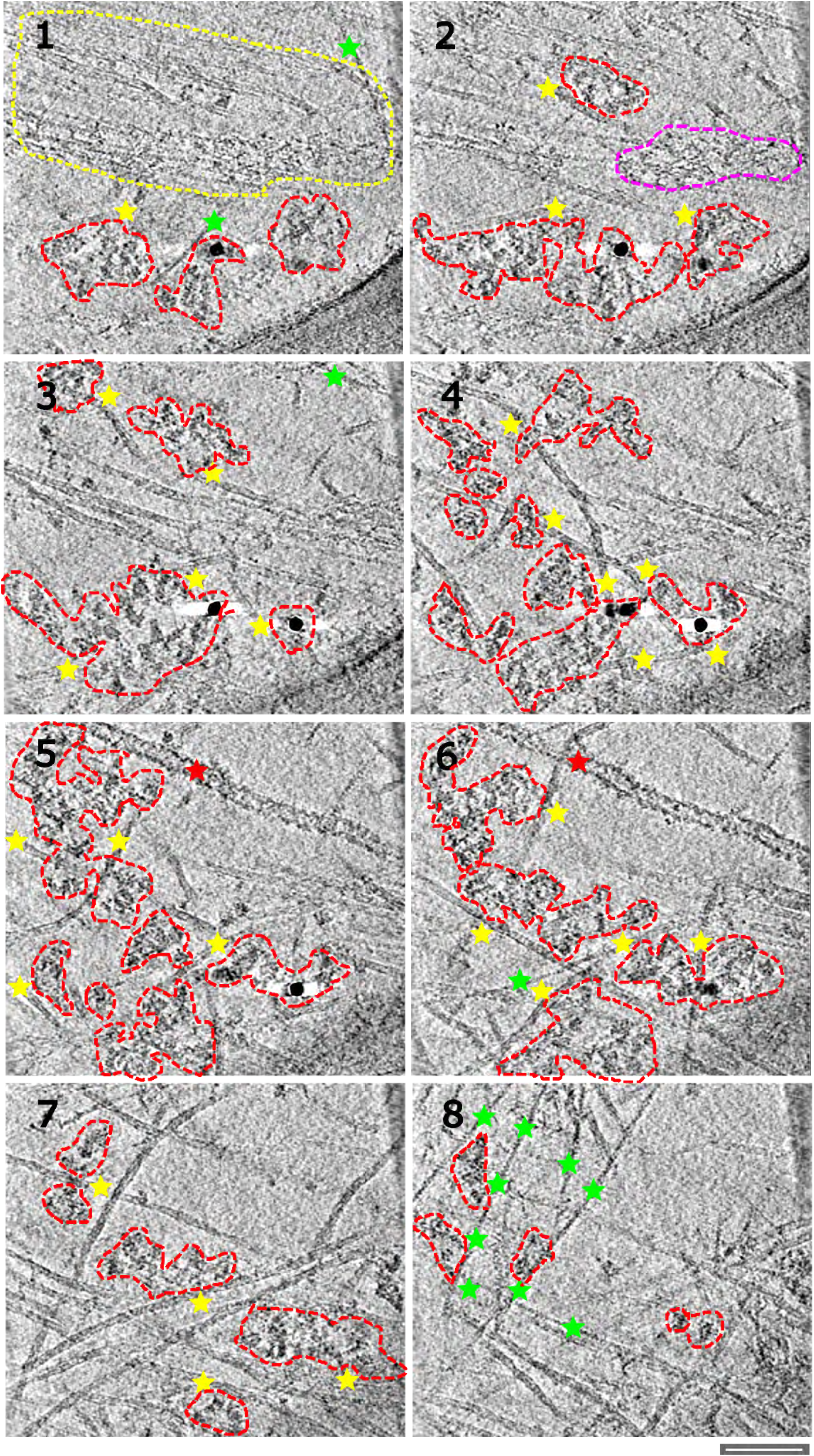
Tomographic slices of the core region from the periphery of the actin bundle. Eight sequential tomographic slices (1–8) from the periphery of the core region are shown, arranged from lower to higher Z positions. Gold particles indicate FA positions, and FA-periphery densities are delineated by red dashed outlines. Magenta dashed outlines indicate protein densities without gold labeling. Yellow dashed outlines indicate the position of the vimentin bundle. Microtubules (red stars), vimentin intermediate filaments (yellow stars), and actin filaments (green stars) are observed either contacting or positioned adjacent to these FA-periphery densities. Scale bar, 100 nm.

A three-dimensional segmentation of this tomogram (**Video 1**) clarified the key features: The top view showed morphologically diverse and discontinuous FA-peripheral densities (**Fig. 7a,b**). An irregular network of vimentin filaments contacted the densities laterally but also penetrated them. A microtubule formed extended contacts with one side of the FA-peripheral protein cluster, and a sparse F-actin layer capped the top surface of the FA-peripheral proteins (**Fig. 7c-d)**.

**Figure 7.**
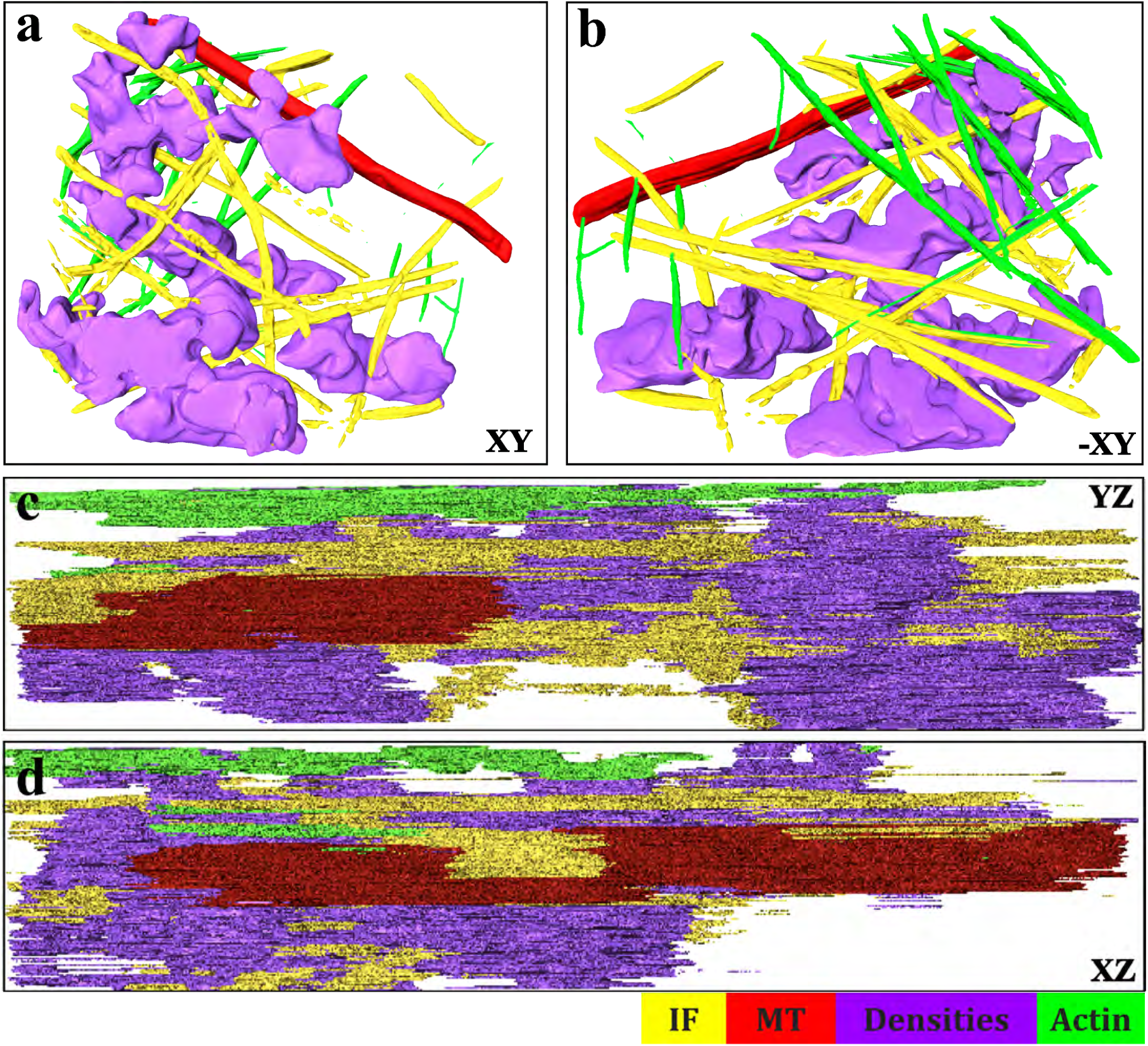
3D segmentation of a tomogram from the periphery of the actin bundle in the core region. Three-dimensional segmentation of a representative tomogram acquired from the periphery of the actin bundle in the core region. FA-periphery densities (magenta), microtubules (red), actin filaments (green), and intermediate filaments (yellow) were segmented in Avizo and visualized as smoothed surface renderings. Panels (a) and (b) show the segmented structures in the XY view, corresponding to the bottom and top views of the same surface rendering, respectively. Panels (c) and (d) show orthogonal views of the segmented volume in the YZ and XZ planes, respectively. These views illustrate the three-dimensional spatial organization of FA-periphery densities relative to surrounding cytoskeletal filaments. Color code is indicated below.

Axis projections of the segmented tomogram further resolved their vertical organization (**Supplementary Fig. 8a**). The XY projections showed the approximately perpendicular orientation of the microtubule relative to the actin fibers. The vimentin filaments did not show preferred orientations in this projection and intercepted the FA-peripheral protein densities from different directions (**Supplementary Fig. 8c)**. The connected protein densities in this location varied significantly in diameter. The YZ and XZ projections (**Supplementary Figure 8d-e**) showed that the sparse F-actin layer occupied only the highest vertical position (corresponding to apparent Z-heights of about 90–100 nm; these heights may be overestimated as they were calculated without tilt-offset corrections; see **Methods**). The ∼8 nm thickness of this F-actin projection is consistent with a defined single-layer arrangement. The microtubule positioned at an intermediate height (∼45–65 nm), whereas vimentin filaments occurred at all Z-heights except for the top actin layer. The thinnest FA-peripheral protein clusters reached about 50 nm in Z-height, which is comparable to the ∼40 nm FA-core thickness reported previously^6^.

However, these projections also revealed that not all the protein densities anchored to the substratum (**Supplementary Fig. 8e**). Collectively, this analysis revealed a three-dimensional architecture where morphologically diverse protein clusters associated in a stratified manner with the different types of cytoskeleton.

### Spatial features characteristic of FA-peripheral densities at the periphery of the actin tip

We next investigated the spatial arrangement in the immunogold membrane-extraced tomograms from the tip region. Cryo-ET analysis of these samples revealed an asymmetric distribution of gold particles (**Fig. 8a**). In the top view in **Fig. 8a**, the gold labels located along the lower-right side of the actin-bundle tip and extended along the right margin of the bundle. Sequential Z-slices of this tomogram revealed several structural features. A dense cluster of FA-peripheral assemblies (**Fig. 8b, 1-2;** red dashed outlines), surrounded by multiple gold particles and accompanied by loosely aligned actin filaments that exhibited a preferential orientation toward the right side of the actin bundle (**Fig. 8b, 1,3;** green stars). Vimentin filaments extended directly into this FA-peripheral protein cluster (**Fig. 8b, 1-2;** yellow star), akin to the vimentin networks that penetrated FA-peripheral densities at the periphery of the actin bundle core. Several F-actin filaments emerging from the actin bundle tip were also observed contacting the FA-peripheral assemblies associated with gold particles(**Fig. 8b, c**, green star; **Supplementary Fig. 9**).

**Figure 8.**
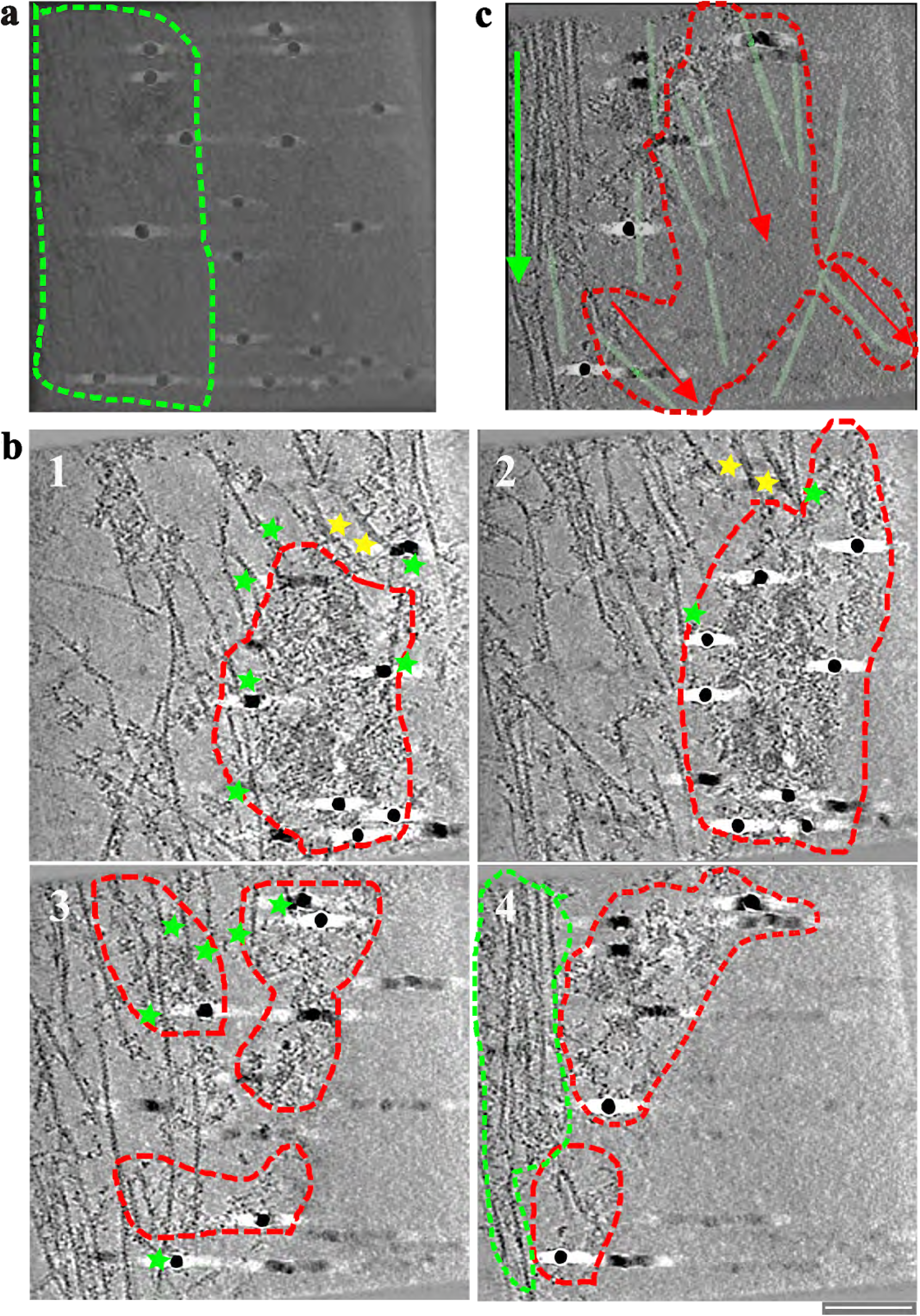
Orientation of actin filaments at the periphery of the tip region. (a) Top-view image of a tomogram volume acquired at the periphery of the tip region, in which multiple gold particles mark FA positions. Green dashed outlines indicate the underlying actin bundle. (b) Sequential tomographic slices arranged from lower to higher Z positions, as indicated by numbers 1–4. Gold particles mark FA positions, and red dashed outlines delineate the corresponding FA densities. Green dashed outlines indicate the actin bundle at the tip region. Yellow stars mark vimentin filaments, whereas green stars mark actin filaments contacting the FA densities. Scale bar, 100 nm. (c) Top-view image illustrating the angular differences between the principal orientation of the actin bundle (green arrow) and the orientations of individual actin filaments associated with gold particles (red arrows) at the periphery. Green dashed lines delineate the actin-bundle boundary at the tip region, and red dashed outlines highlight actin filaments projecting toward the right side of the bundle. To facilitate the visualisation of actin filaments in the periphery, we spatially aligned ∼150 nm long 5 Å F-actin density maps (pale green) with the filament densities associated with gold particles.

Notably, although both F-actin and vimentin directly contact FA-peripheral densities in the tip and core regions, their spatial organization differs markedly: vimentin forms network-like cages around FA-peripheral densities in the core region, whereas such encapsulating vimentin meshes are absent at the tip. Instead, the FA-peripheral densities at the tip are primarily coordinated by F-actin filaments, which surround or directly contact these assemblies.

### Direct and Indirect Vimentin Connections Integrate Vimentins into the Peripheral FA Architecture

Our analysis showed that FA-peripheral protein clusters are located within a heterogeneous cytoskeletal network, in which vimentin filaments either penetrate deeply into the FA-peripheral protein densities or lie closely along their surfaces. To further clarify how vimentin engages these densities in the latter case, we examined additional tomograms taken from the tip region. In several of these datasets, thin linear linker densities were observed bridging vimentin filaments and nearby F-actin filaments at the periphery of FA-peripheral densities. These actin filaments extended directly into the FA-peripheral densities (**Fig. 9a–c)**. The linkers connecting actin and vimentin varied widely in their length (Examples for linker length of 20 nm, 33 nm, and 117 nm are given in **Fig. 9a,b,c,** respectively). The diversity in length, together with the morphology and flexible appearance of this linker, was consistent with plectin, a cytolinker that binds both vimentin and F-actin through its intermediate filament–binding and actin-binding domains^31^.

**Figure 9.**
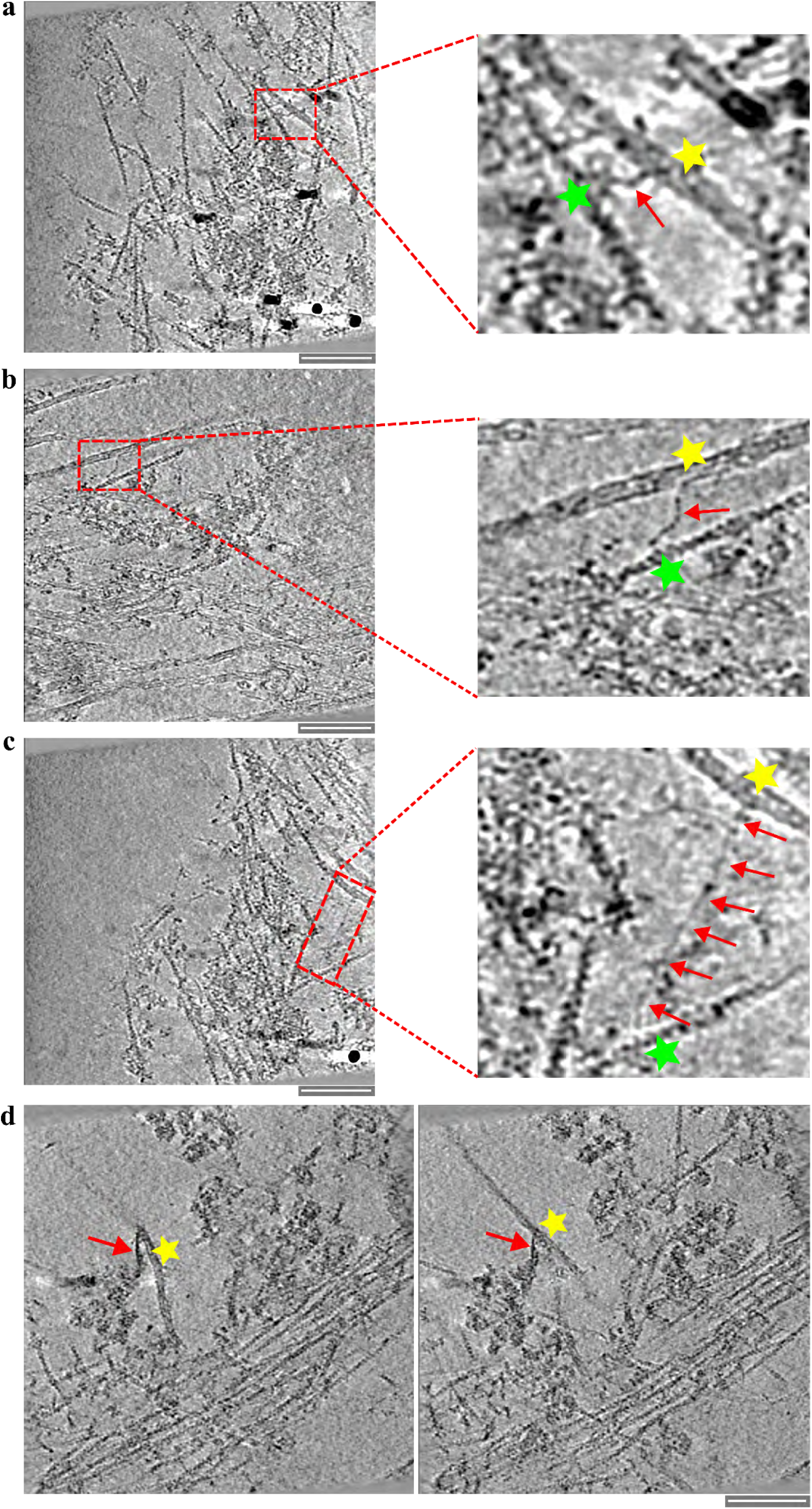
Tomographic slices showing linear connector filaments bridging vimentin, actin, and FA-periphery densities. (a–c) Tomographic slices showing representative examples of linear connector filaments linking vimentin filaments to individual actin filaments. Red dashed boxes indicate the regions displayed as zoomed-in views on the right, where green stars mark the connector filaments of varying lengths. (d) Two tomographic slices (from lower to higher Z positions) from a representative tomogram at the periphery of the tip region, showing a linear density (red arrow) between a vimentin filament and the FA-periphery densities across consecutive slices. Red stars indicate the connecting protein segment. More comprehensive series of tomographic slices spanning low to high Z positions for the same sampling sites are shown in Supplementary Fig. 8. Scale bar, 100 nm.

In addition to these indirect linkages, tomograms also revealed direct contacts between vimentin filaments and FA-peripheral densities. As shown in a representative tomogram in **Fig. 9d (**red arrow), we observed a short linear density extending from a vimentin filament to the surface of the FA-peripheral densities, indicating a direct anchorage of vimentin to the adhesion complex (**Supplementary Fig. 10)**. Taken together, these findings provide an *in situ* observation for protein linkers, presumably plectin isoforms, that tether vimentin fibres to either FA-peripheral densities or to actin filaments.

## DISCUSSION

Biochemical and super-resolution studies have established that the FA core adopts a vertically stratified architecture^21,6,12^. Although cryo-ET was used to structurally characterize specific actin-associated particles^22,23^, the mesoscale spatial organization of the FA-cytoskeleton landscape has not yet been visualised *in situ*. To address this gap, we developed a semi-correlative workflow to map the FA environment at the leading edge of human fibroblasts, based on the analysis of more than 300 tomograms.

Our initial tomogram acquisition was guided by vinculin-targeted fluorescence-labeled FA plaques and performed on intact cells. Vinculin, which associates with talin and paxillin, is a core component of FAs and thus a highly specific marker of these sites^32,22^. To enhance image contrast and resolution, we devised a protocol that removes cell membranes and non-adherent cell content without disrupting FA-peripheral structures. These thinner samples, labelled with anti-vinculin immunogold, revealed canonical FA-core densities beneath the actin bundles. These densities were not visible in whole-cell tomograms likely due to poor contrast caused by the overlying thick actin bundles. These observations may explain why a previous cryo-ET study did not capture FA-core densities^22^.

Using these preparations, we were able to support the identification of the intermediate filaments as vimentin by subtomogram averaging. Indeed, vimentin is known to be the major intermediate filaments in fibroblasts. Keratin is very unlikely to be expressed in these cells, and keratin filaments typically consist of eight protofilaments^33^. However, we cannot rule out that the filaments we observed also contain nestin, which forms heterofilaments with vimentin, but does not assemble into filaments on its own^34^. Vimentin is particularly relevant to FA biology because it directly associates with integrin–positive adhesions, extends coordinately with adhesion growth, and supports the formation of larger, mechanically reinforced FAs^18^. Our 12.96 Å reconstruction resolves the protofibril spacing with greater clarity than the ∼20 Å cryo-ET map previously reported^15^. Our map revealed the characteristic five-protofibril cross-section, expected helical rise and twist, and closely fit to the atomic model of mature vimentin (PDB 8RVE). This structural match further indicated that the filaments observed *in situ* correspond to mature vimentin. Indeed, mature vimentin filaments, but not precursors, are known to regulate FA dynamics, force propagation, and adhesion turnover^19,17,35^.

Collectively, our cryo-ET analysis offers a comprehensive *in situ* view of the adhesion landscape, extending from the substratum to the overlying FA components at a height of ∼100 nm (**Fig. 10**). At the actin core region, actin bundles appeared as densely packed, uniformly aligned filaments with large, continuous FA-core densities positioned beneath them, in line with high-tension adhesions from previous super-resolution studies^6,12^. The periphery flanking the actin core displayed a stratified organization: ∼15 nm–thick basal layer of roughly parallel vimentin filaments lay beneath an intersecting vimentin mesh occupying the region ∼15–90 nm above the substratum. These vimentin filaments surround and occasionally penetrate FA-peripheral protein clusters. These clusters were morphologically diverse, and sometimes formed contiguous groups. They occupied the entire vertical span (Z = 0–100 nm), but were largest between 20 and 80 nm, as not all connected to the substratum or top layers. A sparse F-actin layer, loosely perpendicular to the central actin bundle, topped the tomographic structure at heights of ∼90–100 nm, with occasional filaments extending downward into FA-peripheral densities. A single microtubule was typically observed in each tomogram at intermediate heights (Z = 45–65 nm), oriented parallel to the actin bundle.

**Figure 10.**
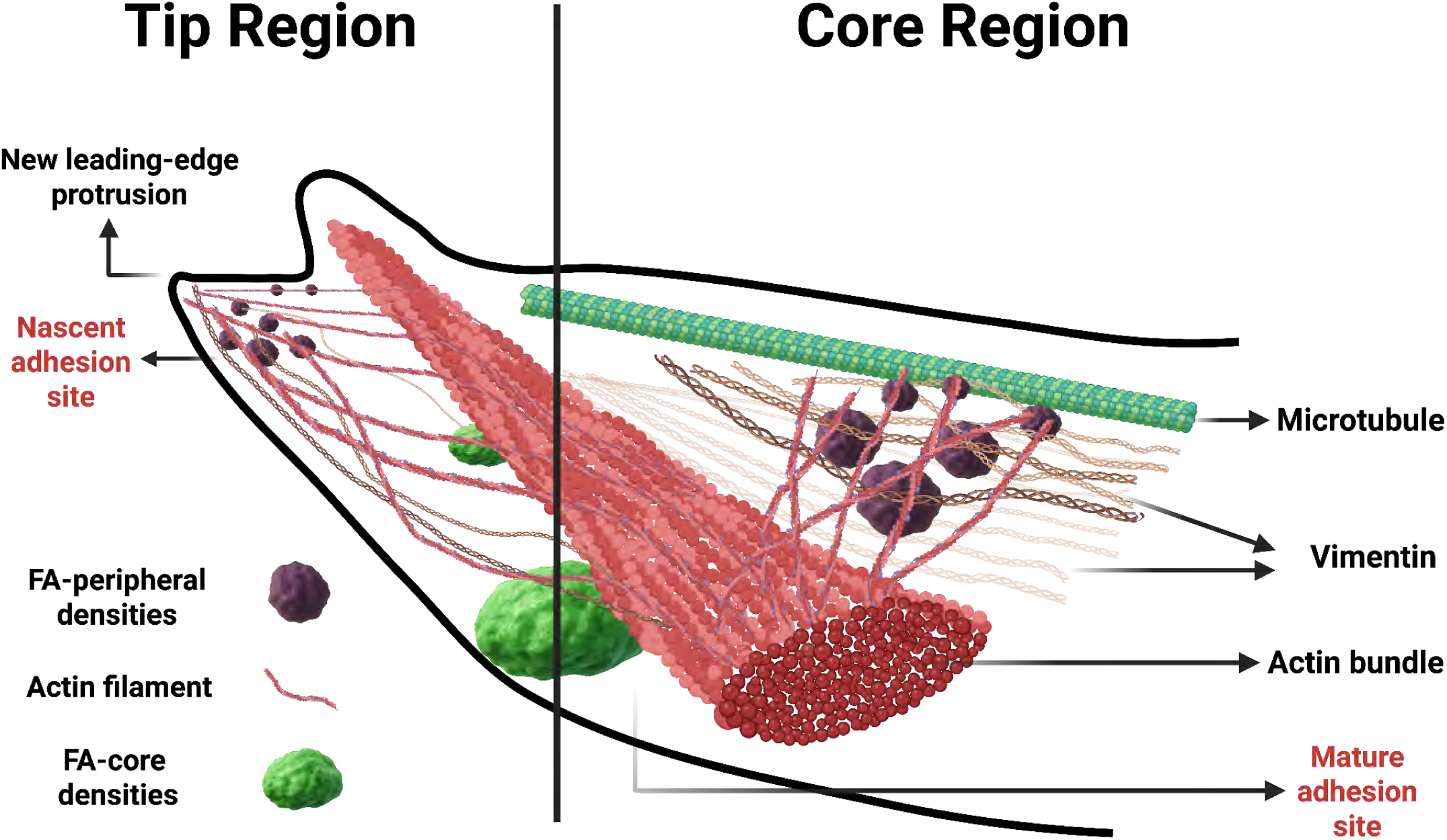
Schematic model of distinct FA architectures at the tip and core regions of actin bundles. This schematic summarizes the in situ architectures of FA-peripheral and FA-core densities identified at the tip and core regions along actin bundles. At the tip region (left), corresponding to nascent adhesion sites at the leading edge, FA-peripheral densities accumulate along the lateral margins of the actin bundle. These assemblies adopt an actin-dominated architecture in which FA-peripheral densities interact with actin filaments and vimentin within a single cytoskeletal layer. In this region, vimentin contacts FA-peripheral densities but does not form ordered cage-like meshes. At the core region (right), corresponding to mature adhesion sites, FA assemblies exhibit a vertically stratified architecture. FA-core densities are positioned beneath the actin bundle, whereas FA-peripheral densities are embedded within a multilayered cytoskeletal system. In this configuration, vimentin forms ordered arrays and meshworks that penetrate or abut FA-peripheral densities, a microtubule occupies an intermediate layer, and a sparse actin sheet caps the uppermost layer. Together, this model reveals that FA assemblies adopt two spatially and structurally distinct architectures along actin bundles: a core architecture located beneath the bundle and a peripheral architecture distributed along its margins. FA-peripheral densities engage the cytoskeleton through distinct structural modes, with actin filaments inserting into or attaching to the densities, while vimentin interacts via insertion, actin-mediated indirect attachment, or direct contact.

In contrast, the actin tip region featured thinner, less dense actin bundles without large FA-core densities observed directly beneath. The periphery of the actin bundle tip contained sparse, irregular F-actin and vimentin filaments interspersed with discontinuous FA-peripheral clusters. Vimentin and actin filaments frequently surrounded or penetrated these protein clusters. Plectin-like linkers connected vimentin either to the protein clusters or to actin filaments. Many actin filaments terminating in FA-peripheral densities could be traced back to the actin bundle core. The reduced coupling of these FA-peripheral clusters to actin and vimentin at the lateral periphery of the tip region, and their position within leading-edge protrusions, suggested that this landscape represents nascent, low-tension adhesions^19^ (**Fig. 10**).

In summary, our study provides an *in situ* structural framework for how FA-peripheral assemblies engage in adhesion formation across different phases (**Fig. 10**). Beneath the actin bundle core region, large continuous FA protein clusters form large, high-tension adhesions associated laterally reinforced by thick vimentin bundles under shear stress^18^. FA clusters peripheral to the actin core are embedded in a dense but more loosely organized network formed by vimentin, actin filaments and microtubules. The peripheral network and its engagement with FA-peripheral clusters weakens toward the tip of the actin bundle, suggesting that these protein clusters represent nascent adhesions poised for leading-edge protrusion^19^. The presence of direct and tethered contacts between vimentin, actin and FA-peripheral clusters in this area suggests their spatial and mechanical coordination during adhesion growth. Our findings extend classical lamellipodial FA models to peripheral adhesion zones and provide a mechanistic basis for how dispersed FA assemblies integrate multiple cytoskeletal systems to sustain force transmission and drive FA maturation during cell migration.

## METHODS

### Cell Culture

Fibroblasts were isolated from healthy newborn males during circumcision procedures and established with informed consent and under protocols approved by the respective institutional review boards. Cells were routinely tested for mycoplasma contamination and maintained in Dulbecco’s Modified Eagle’s Medium (DMEM, GlutaMAX; Thermo Fisher Scientific, #31966021) supplemented with 10% (v/v) fetal bovine serum (Thermo Fisher Scientific, #26140087). Cultures were incubated at 37 °C in a humidified atmosphere containing 5% CO₂.

### Grid Preparation and Cell Seeding

Gold 200-mesh holey carbon finder grids with a 2 nm additional carbon support film (R1.2/1.3, LF; Quantifoil, LFH2100AR1.3) were glow-discharged using a PELCO easiGlow™ system for 30 seconds. Pretreated grids were mounted onto glass-bottom dishes (Willco, #HBST-3512) using custom-designed stencils as previously described^36^. Grids were exposed to UV light for 30 min in a biosafety hood and subsequently coated with 0.1 mg/mL fibronectin (Fisher Scientific, #11564466) for 2–4 h at room temperature (RT). After coating, grids were washed three times with PBS (Gibco, #14190094). Freshly trypsinated fibroblasts were seeded directly onto the grids and incubated overnight at 37 °C with 5% CO₂ to allow cell adhesion. Following incubation, cells were subjected to different treatments.

### Immunofluorescence Staining on Grids

Cells cultured on EM grids were fixed with 4% EM-grade PFA (Fisher Scientific, #50-980-487) for 10 min under a chemical hood, followed by permeabilization with 0.1% Triton X-100 in PBS (PBST) for 30 min. Samples were blocked with 1% BSA (Merck Millipore, #A3294-50G) in PBST for 1 h at RT, then incubated overnight at 4 °C with Alexa Fluor 488-conjugated anti-Vinculin antibody (Abcam, #ab196454; 1:100 dilution). Between each step, grids were washed three times with PBS. Fluorescence imaging was performed using a Leica Stellaris Falcon_iac confocal microscope. Cells exhibiting distinct FA plaques adjacent to finder grid markers were identified and documented to enable precise correlation and relocation of the same regions during subsequent Cryo-ET.

### Cell Membrane Extraction and Gold Immunolabeling of FAs

Cell membrane extraction was performed as previously described^37^. Briefly, cells were treated for 1 min with cytoskeleton buffer [10 mM MES (pH 6.2), 150 mM NaCl, 5 mM EGTA, 5 mM glucose, 5 mM MgCl₂] containing 0.75% Triton X-100 and 0.25% glutaraldehyde to dissolve the plasma membrane. Samples were then fixed with 2% glutaraldehyde in the cytoskeleton buffer for 15 min at RT before vitrification or subsequent immunogold labeling.

To quench residual aldehyde groups, grids were incubated with 0.05 M glycine in the cytoskeleton buffer for 15 min, followed by blocking with 5% BSA in the same buffer for another 15 min. Between each step, grids were washed three times (3 min per wash) with PBS. Grids were then incubated overnight at 4 °C with rabbit recombinant monoclonal anti-Vinculin antibody (Abcam, #ab207440; 1:800 dilution) in 1% BSA, followed by incubation with anti-rabbit IgG (H&L) conjugated to 15 nm gold particles (AURION, Lot GAR-90723/1; 1:20 dilution) overnight at 4 °C. After six washes (5 min each) with blocking buffer, grids were vitrified and stored in liquid nitrogen until imaging.

### Data Collection of Differently Treated Samples

Tilt-series were acquired from distinct cellular regions according to the sample preparation from different workflows. For immunofluorescence-stained samples, areas containing vinculin-positive plaques were selected for imaging. For membrane-extracted samples, lateral actin bundle or its tip body region were chosen. In membrane-extracted samples with immunogold labeling, regions containing gold particles near actin bundle ends were selected.

Tilt-series were collected on a Titan Krios transmission electron microscope (Thermo Fisher Scientific) operated at 300 keV and equipped with a Falcon 4i direct electron detector. The microscope was operated with a 50-μm C2 aperture and an energy filter slit width of 10–20 eV. In total, over 100 tilt-series were recorded at magnifications of ×105,000 corresponding to a pixel size of 1.19 Å using Tomography 5 software (Thermo Fisher Scientific). Multi-frame movies were acquired over a tilt range of −60° to +60° in 2° increments, with a frame rate of 7 frames per second and an electron flux of ∼2.0 e⁻/pixel/s. The cumulative dose was ∼120 e⁻/Å², and the nominal defocus range was −2 to −7 μm.

### Tomogram Reconstruction and Preprocessing

Tilt-series stacks were aligned and reconstructed in IMOD^38^ after 3× binning for tomogram screening and region-of-interest selection. Selected multi-frame movies of tiltseries were imported into RELION 5.0 for motion correction, contrast transfer function (CTF) estimation, tiltseries alignment with IMOD within RELION 5 and tomogram reconstruction, with or without denoising depending on the analysis purpose. Tomograms were binned to appropriate pixel sizes for subsequent processing: reconstructions with denoising were binned to 10 Å/pixel for visualization and filament picking in Dynamo, while non-denoised tomograms were binned to 4 Å/pixel for high-resolution subtomogram average.

### Subtomogram Averaging and Structural Determination of Intermediate Filaments

To investigate the structural organization of intermediate filaments, tilt series were acquired from regions flanking actin bundle protrusion ends, specifically along both sides of the bundle within the lamellar region, at 300 keV using a Falcon 4i detector and a 10 eV energy filter slit, with a nominal defocus of −3 μm.

Tilt-series stacks were first reconstructed in IMOD to select tomograms enriched in intermediate filaments. The selected stacks were then imported into RELION 5.0 for frame-based motion correction, CTF estimation, tomogram reconstruction, and denoising (for the picking step). The resulting denoised tomograms were used in Dynamo for manual tracing of intermediate filaments, while the non-denoised tomograms were used for subtomogram extraction.

Two sets of extraction parameters were applied in Dynamo: subtomograms were picked every 60 Å with a 380 Å box, yielding 37,582 subtomograms, and every 160 Å with a 500 Å box, yielding 12,596 subtomograms. The extracted subtomograms were then processed using the Actin Polarity Toolbox (APT)^27^. APT was applied to both datasets at projection thicknesses of 130 Å and 330 Å respectively, to generate 2D projection images, which were subsequently subjected to several rounds of unsupervised 2D classification in RELION 5.0 (**Supplementary Fig. 6**). From the 60 Å / 380 Å dataset, high-quality 2D class averages were transferred to cryoSPARC for Homogeneous and Helical Refinement. During the helical refinement, we performed a helical symmetry search in CryoSparc around the reported values for the helical rise of 42 Å and twist of 72°^15^. A grid search was performed over a rise range of 30–55 Å and a twist range of 60–80°, starting from initial estimates of 42 Å rise (Δz, Å) and 72° twist (φ, degrees). Candidate symmetry parameters were evaluated by applying helical operators to the reconstructed volume and computing the mean squared error (MSE) between symmetry-related densities. Local optima were ranked by MSE, and the final rise and twist values were selected from the lowest-error solution for subsequent helical refinement. The helical rise and twist were determined to be 40 Å and 72°, respectively (**Supplementary Fig. 7b**).

The refined density map was fitted to the vimentin filament model (PDB: 8RVE) in UCSF ChimeraX, yielding a cross-correlation coefficient of 0.86. In parallel, class averages from the 160 Å / 500 Å dataset were analyzed in MATLAB by autocorrelation to determine the axial periodicity and repeat length of the intermediate filaments.

## Acknowledgements

We thank our collaborators at the King Faisal Specialist Hospital and Research Center, particularly Lama Alabdi and Dr. Fowzan S. Alkuraya, for providing the human-derived fibroblasts used in this study. We thank Dr. Nagarajan Kathiresan at the King Abdullah University of Science and Technology (KAUST) Supercomputing Core Lab for his assistance with the IBEX computing environment. We thank Kai Qi (University of Tokyo) for designing the computer-aided design files for the 3D-printed stencils used in this work. Computational resources were provided by the KAUST Supercomputing Laboratory. Experimental support was provided by the Bioscience Core Lab, the ACL Proteomics Lab, and the Imaging and Characterization Core Lab at KAUST. The research reported in this publication was supported by funding from KAUST with the baseline fund to STA, and the KAUST Center of Excellence for Smart Health (KCSH), under award number 5932.

## Author contributions

P.Y. developed the experimental workflow, prepared all samples, performed cryo-ET data acquisition, carried out image processing, segmentation, structural analysis, and generated all figures. L. Z. contributed to the early stages of experimental design, assisted with initial data collection, and provided theoretical guidance and critical discussions when methodological questions arose. A.A. guided and assisted sample preparation, data collection, and data analysis. He contributed to troubleshooting experimental procedures, discussed data interpretation with P.Y., and helped refine the experimental strategy throughout the study. S.T.A. supervised the project, secured funding and resources, guided the scientific direction, coordinated collaborations, and provided critical conceptual input throughout the study. P.Y. and S.T.A. wrote the manuscript, and all authors read and approved it.

---

**Supplementary Figure 1.**
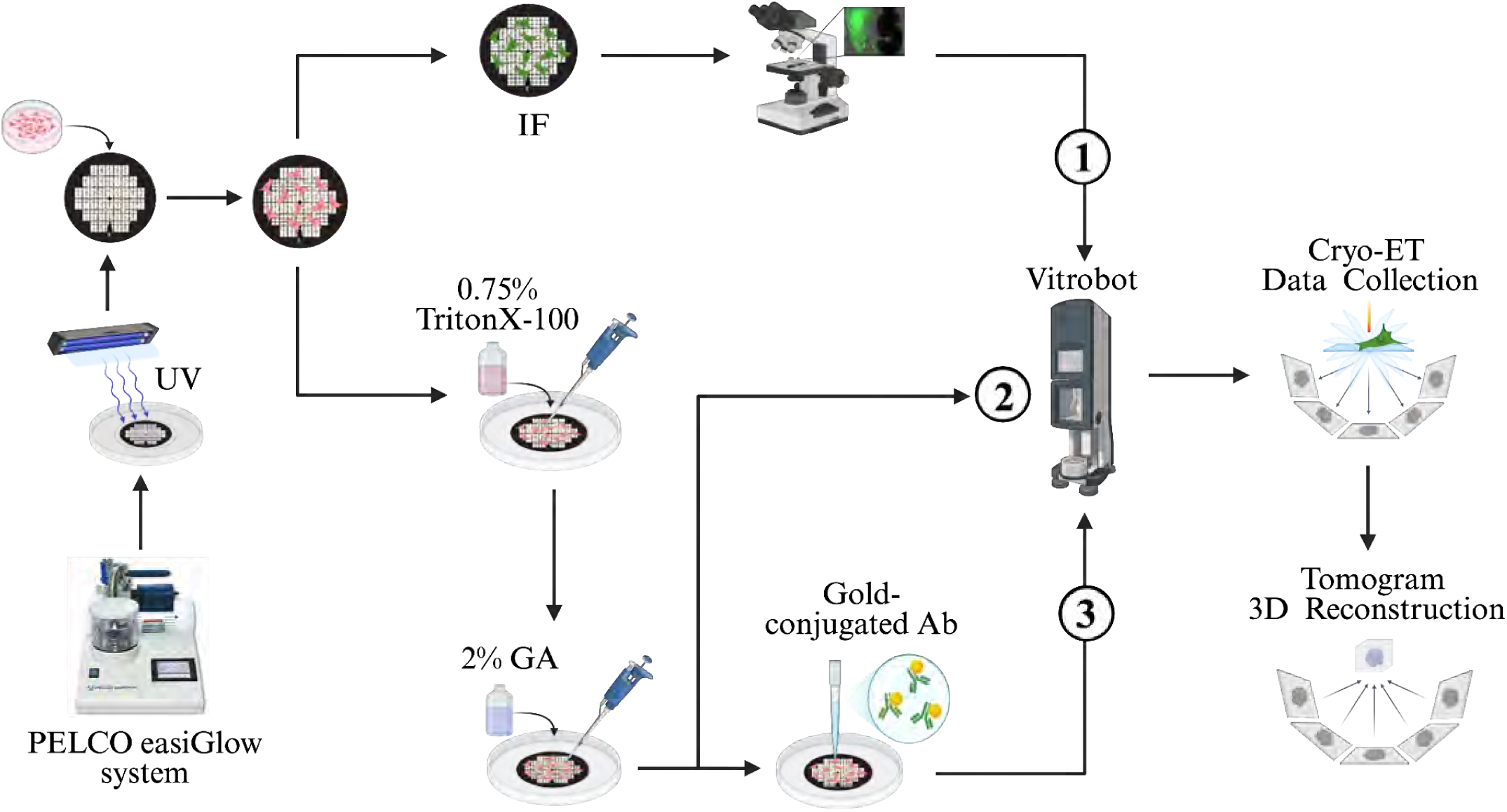
Workflow for cryo-ET sample preparation and correlative analysis. Schematic overview of the workflow for preparing fibroblast grids for cryo-ET. Gold 200-mesh Quantifoil finder grids were glow-discharged using a PELCO easiGlow™ system, UV-treated, coated with fibronectin, and seeded with fibroblasts. In route 1, fibroblasts were fixed with 4% PFA and subjected to immunofluorescence staining for vinculin. In route 2, fibroblasts were treated with 0.75% Triton X-100 to extract the plasma membrane and then fixed with 2% GA. In route 3, gold-conjugated anti-vinculin antibodies were added to route 2 for immunogold labeling. All grids were vitrified using a Vitrobot and stored in liquid nitrogen for cryo-ET data collection and 3D tomogram reconstruction.

**Supplementary Figure 2.**
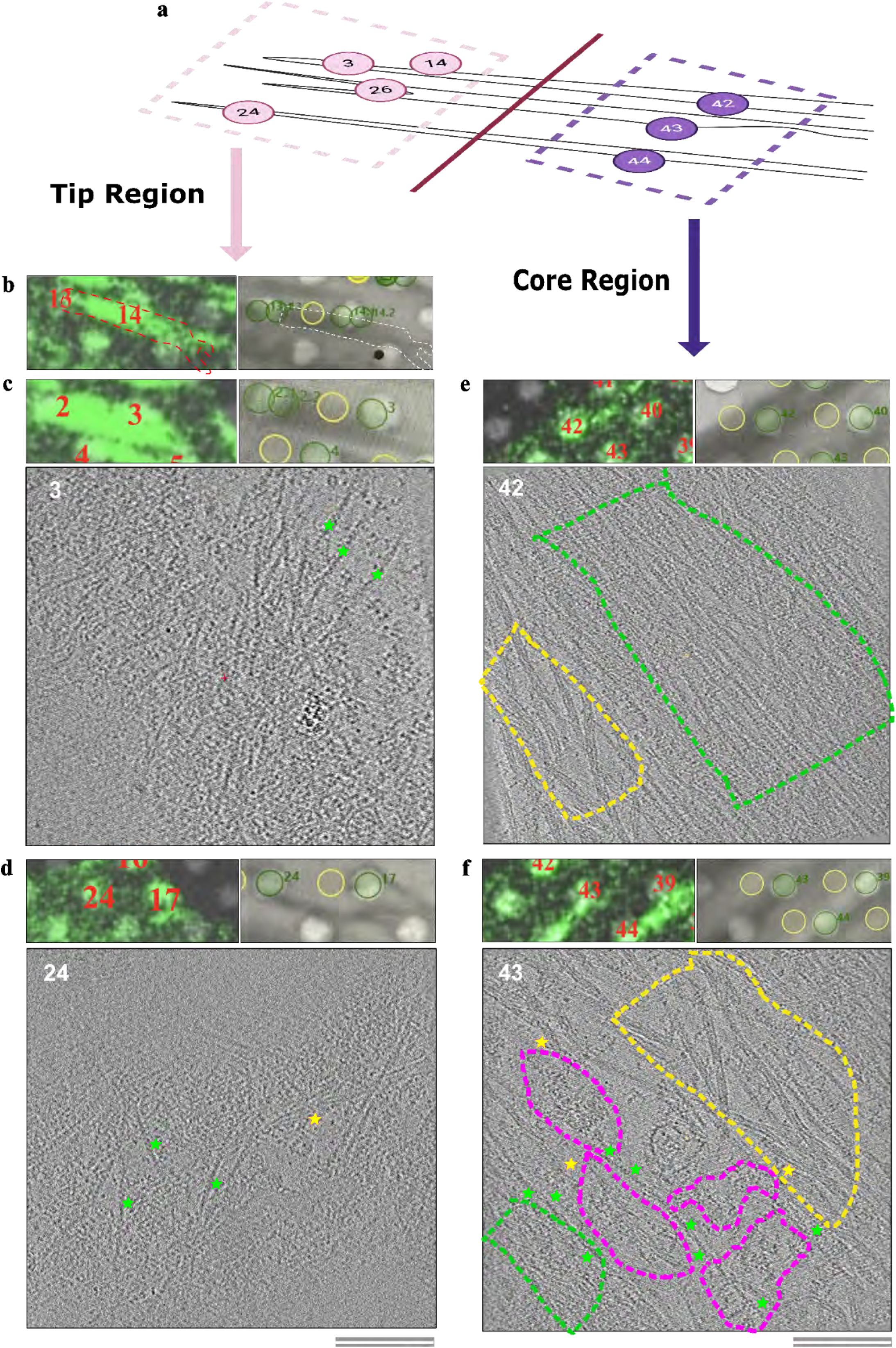
Correlative fluorescence–cryo-ET analysis of FA sites at the tip and core regions. (a) Schematic illustrating cryo-ET sampling positions along a leading-edge actin bundle. The brown line delineates the Tip Region, located near the actin bundle tip, from the Core Region, positioned posterior to the tip. Numbered circles indicate the cryo-EM sampling sites analyzed in this study. (b) Correlative fluorescence image showing vinculin-positive FA sites (left) and the corresponding cryo-EM grid map (right). The extent of the filament bundle identified in the EM grid view (white dashed outline) is projected onto the fluorescence image and indicated by the red dashed outline. (c–f) Representative correlative fluorescence–cryo-ET views from sampling positions 3 (c) and 24 (d), representing the tip region, and positions 42 (e) and 43 (f), representing the core region. For each position, the upper left panel shows the fluorescence localization of FA sites, the upper right panel shows the corresponding cryo-EM grid view with circles indicating selected coordinates, and the lower panel displays a representative tomographic slice from the indicated sampling position. In the tomographic slices (e, f), magenta dashed outlines indicate dispersed protein densities, green dashed outlines denote dense actin-bundle regions, and yellow dashed outlines mark intermediate filament–rich regions. Green stars indicate actin filaments contacting protein densities, and yellow stars indicate intermediate filaments contacting protein densities. In the tip region (c, d), only sparse individual actin (green stars) and intermediate filaments (yellow stars) are observed, whereas in the core region (f), multiple actin and intermediate filaments attach to dispersed small protein densities. Scale bar, 100 nm.

**Supplementary Figure 3.**
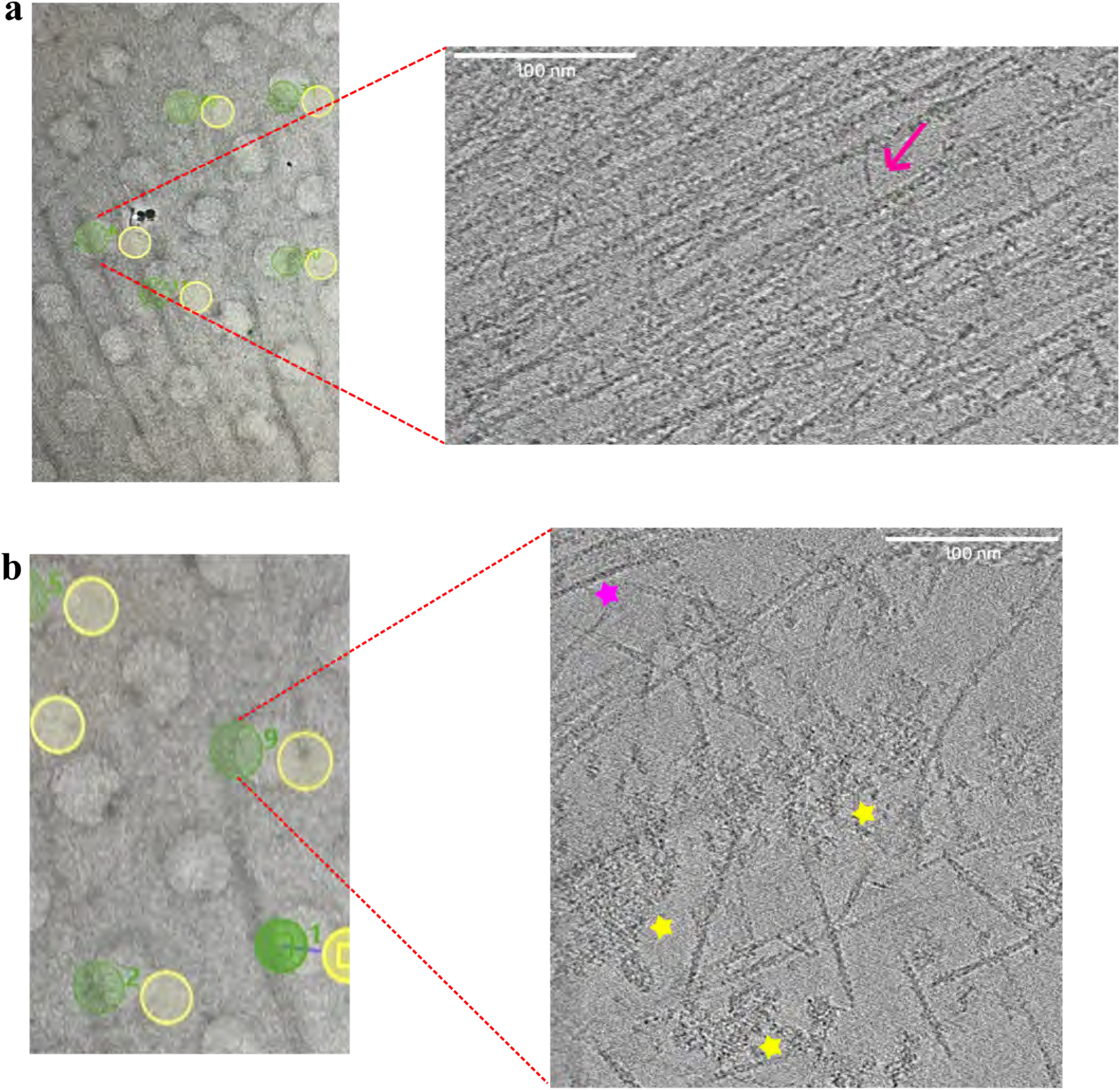
Cryo-electron tomography analysis of actin bundle–associated regions in membrane-extracted fibroblast. (a) Tomographic data collected at the tip region of an actin bundle. The left panel shows the low-magnification TEM view of the grid area, with the position used for data acquisition highlighted (green circles). The right panel displays a representative Z-slice from the tomogram, showing regularly arranged F-actin filaments. Several filamentous densities link adjacent filaments; one is illustrated by an magenta arrow. (b) Tomographic data collected at the lateral region of the actin bundle. The left panel shows the TEM view indicating selected acquisition sites. The right panel shows a tomographic slice revealing a microtubule (the magenta star) and nearby protein clusters (yellow stars) associated with individual actin filaments.

**Supplementary Figure 4.**
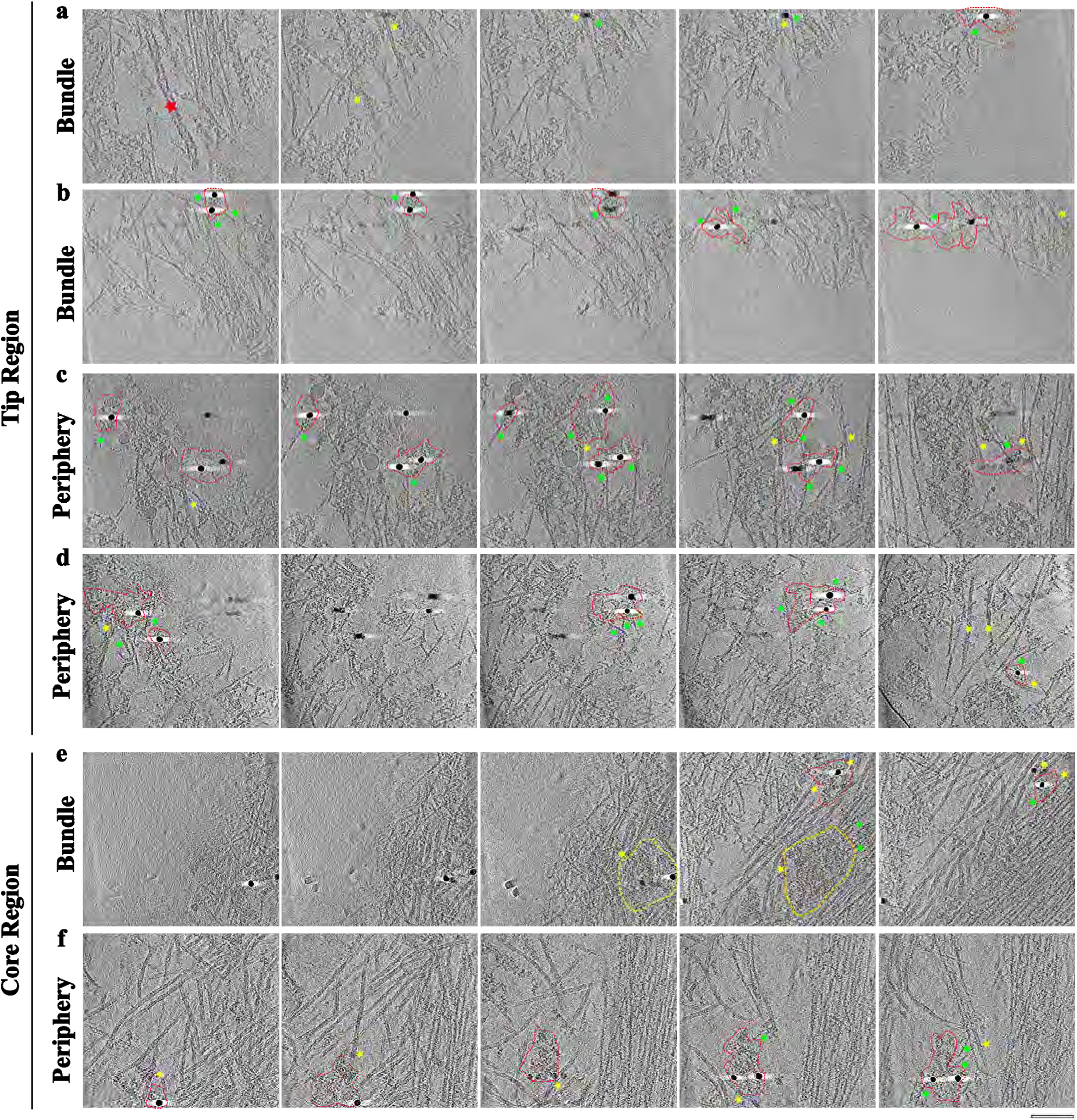
Representative tomographic slices showing FA densities at bundle and periphery regions. Shown are tomographic slices from the tip (a–d) and core (e–f) sampling regions, classified into bundle and periphery zones (Fig. 3). For each region, a series of five sequential Z-slices is displayed from lower to higher positions (left to right). Across regions, filament density and composition differ markedly. Gold particles mark FA locations, where FA-periphery densities (red dashed outlines) or classical FA-core densities (yellow dashed outlines) are observed. MTs (red stars), IFs (yellow stars), and actin filaments (green stars) are observed either contacting or situated near these gold-labeled densities. Scale bar, 100 nm.

**Supplementary Figure 5.**
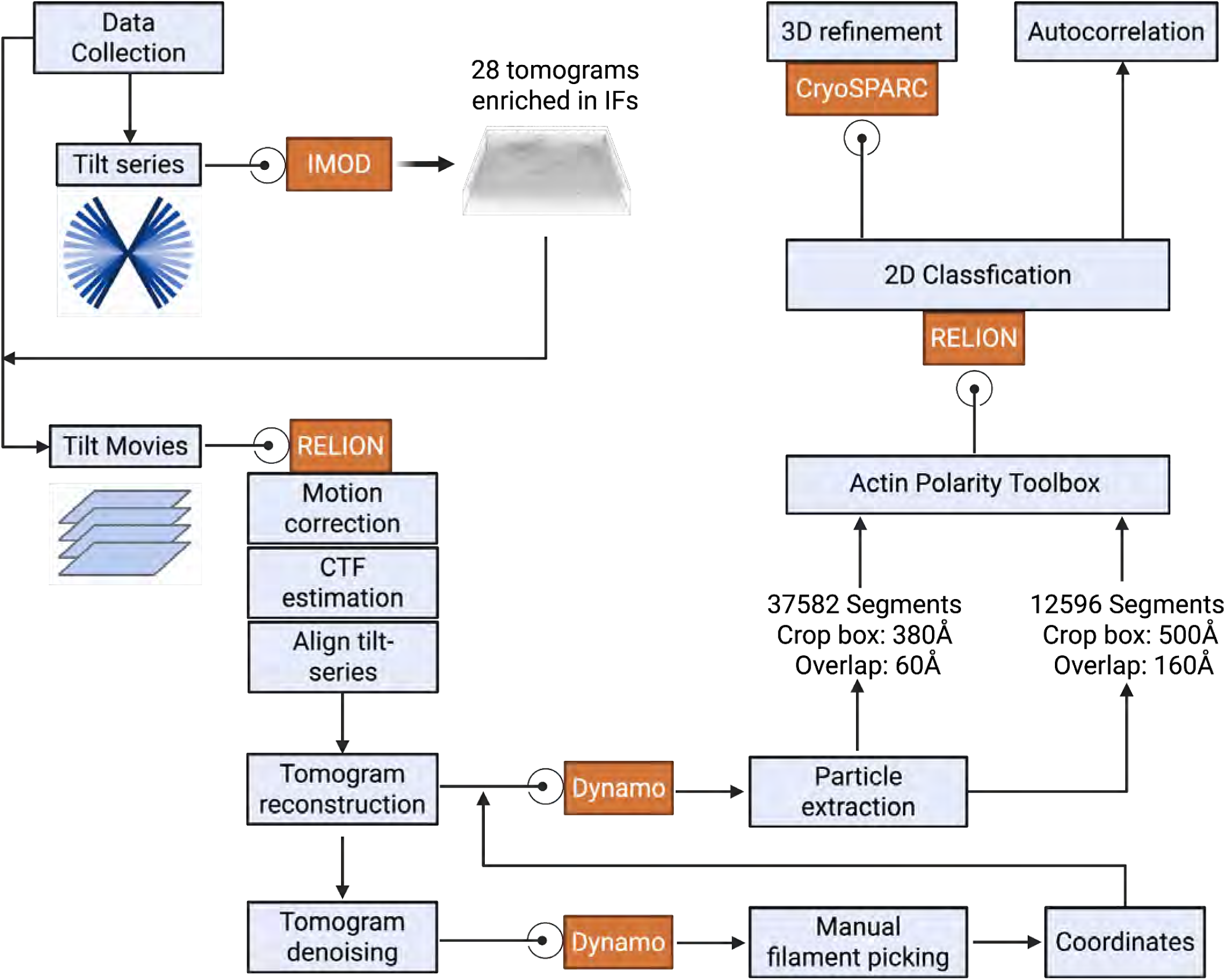
Workflow of structural determination for IFs. Overview of the processing pipeline combining RELION, Dynamo, APT, and CryoSPARC.

**Supplementary Figure 6.**
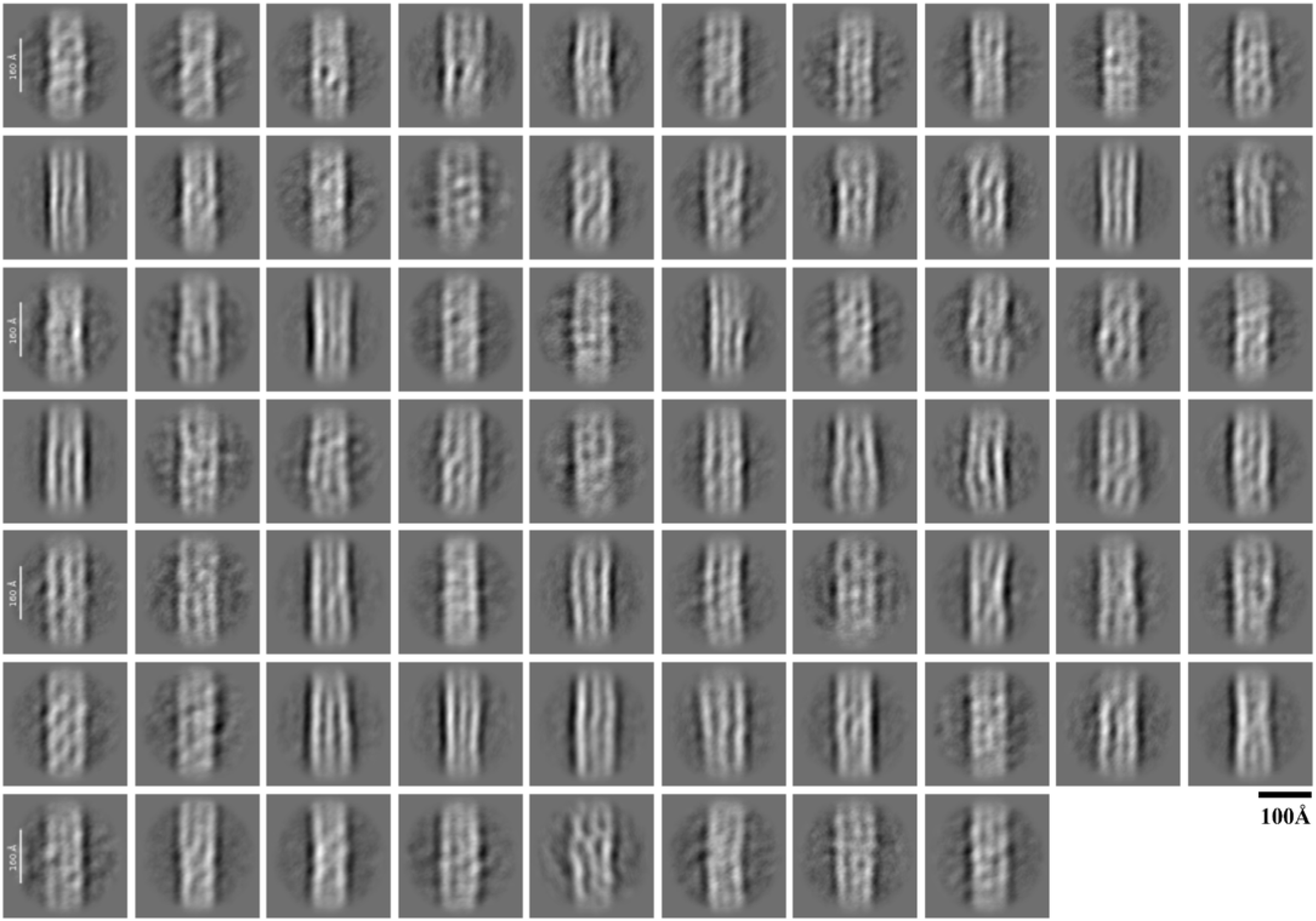
2D classification of intermediate filament segments. A total of 37,582 segments (box size: 38 × 38 nm²) were extracted from tomograms of detergent-treated, gold-labeled fibroblasts. The inter-segment picking distance was set to 60 Å, and the projection thickness to 130 Å. The displayed class averages were generated from ∼6,200 selected particles used for subsequent 3D reconstruction. Scale bar, 100 Å.

**Supplementary Figure 7.**
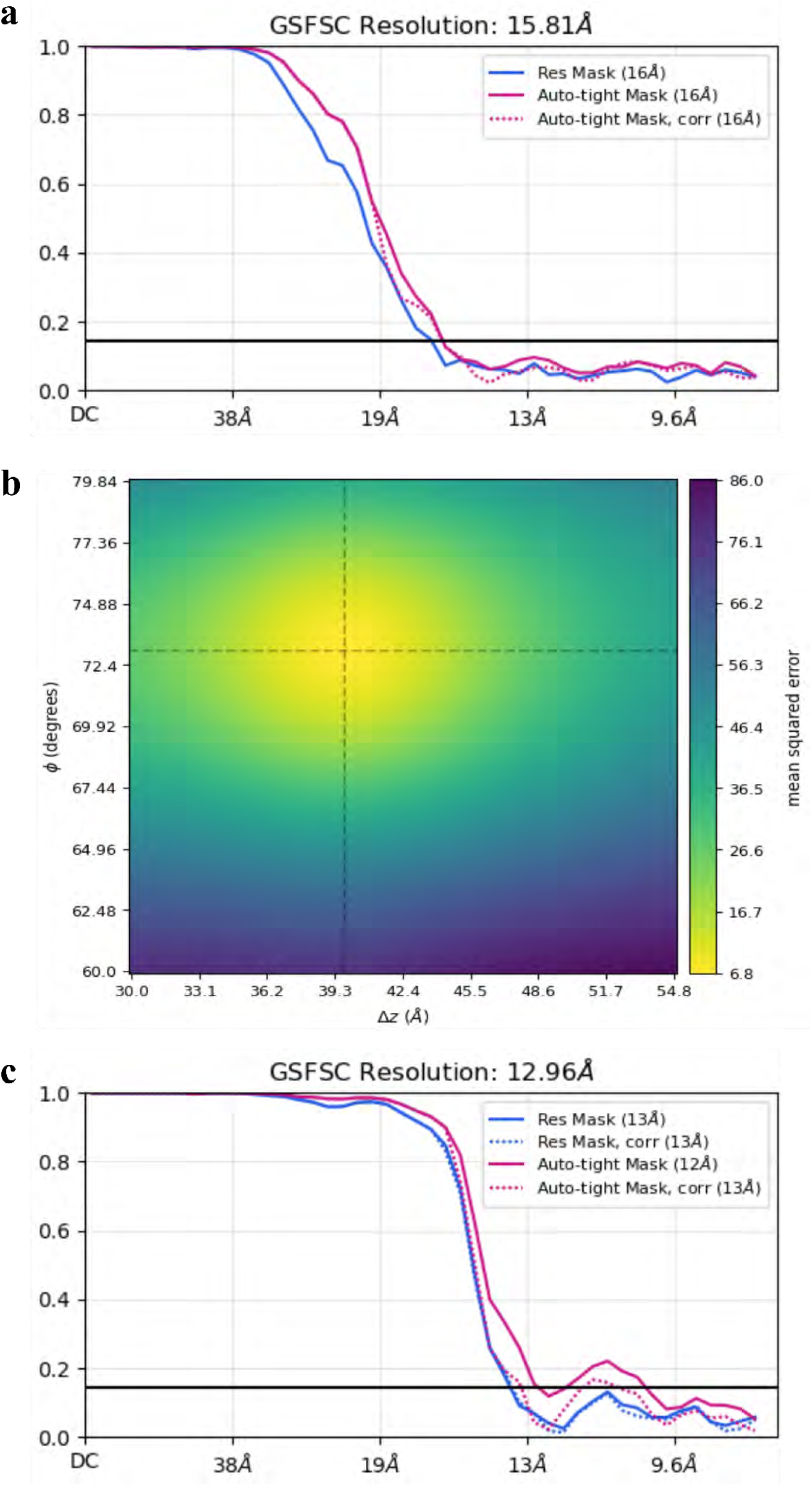
Helical symmetry optimization and resolution assessment of the reconstructed map. (a) Gold-standard Fourier shell correlation (GSFSC; 0.143 criterion) curves of intermediate filament averages obtained from homogeneous refinement, yielding a global resolution of 15.81 Å. (b) Parameter search landscape for helical symmetry determination. The mean squared error is plotted as a function of axial rise (Δz, Å) and helical twist (φ, degrees). The global minimum (yellow region) indicates the optimal helical parameters used for subsequent refinement; dashed lines mark the selected rise and twist values. The helical rise and twist were determined to be 40 Å and 72°, respectively. (c) GSFSC curves of averages obtained from helical refinement after masking optimization, yielding an improved global resolution of 12.96 Å. Solid and dotted lines represent masked and mask-corrected FSC curves, respectively.

**Supplementary Figure 8.**
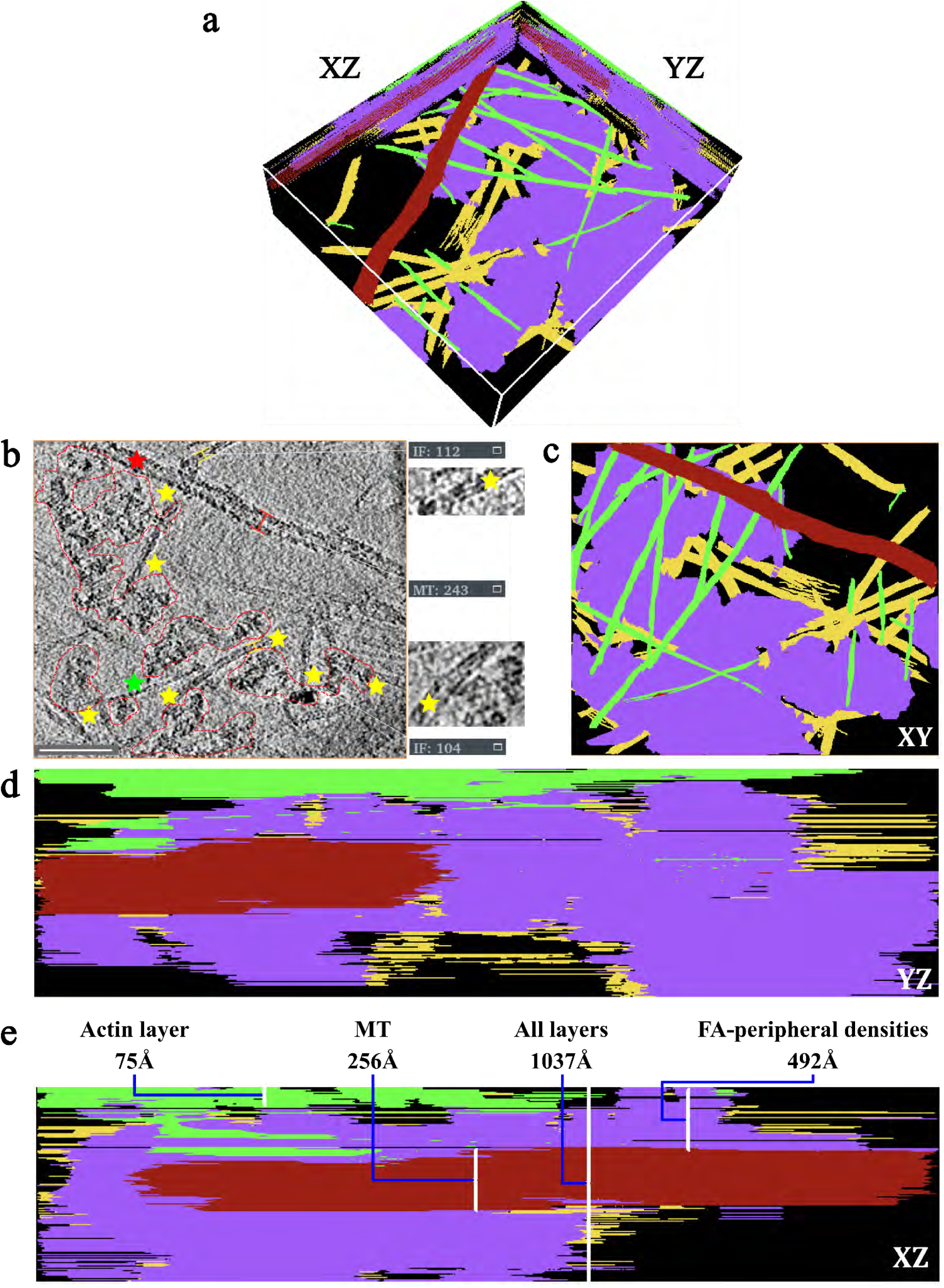
Orthogonal projections of segmented FA-peripheral regions from a tomogram at the periphery of the core region. (a) Combined orthogonal projection generated in Avizo, showing segmented components simultaneously in the XY (c), YZ (d), and XZ (e) orientations. Color scheme: microtubules (MT, red), actin (green), vimentin intermediate filaments (yellow), and FA-peripheral densities (magenta). (b) Representative tomographic slice near the mid-Z plane from the same tomogram, showing MTs (red stars), vimentin filaments (yellow stars), actin filaments (green stars), FA-peripheral densities (red outlines), and gold fiducials within the same field of view. Insets on the right show magnified vimentin segments used for filament diameter measurements. Scale bar, 100 nm. (e) Vertical thickness measurements derived from the XZ projection for each segmented component. “All layers” denotes the combined vertical extent of all segmented structures; “MT”, the vertical thickness of the microtubule; “FA-peripheral densities”, the vertical thickness of FA-peripheral densities; and “Actin layer”, the vertical thickness of the actin projection. All values are reported in ångströms (Å). Color scheme as in (a).

**Supplementary Figure 9.**
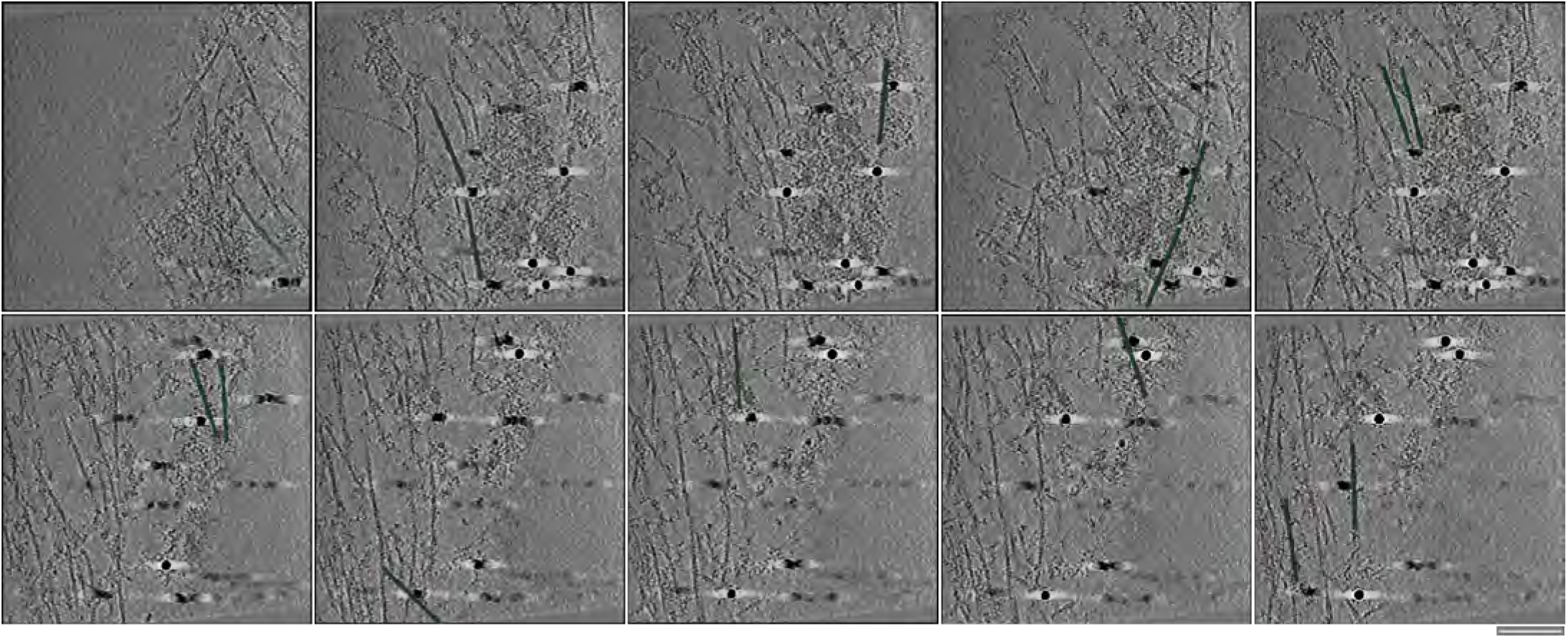
Localization of actin filaments at the periphery of the tip region. Tomographic slices illustrating the overlap between actin filaments and actin densities adjacent to gold particles. A 5 Å F-actin density map (green, 20% transparency) was overlaid onto the corresponding filament densities in the raw tomogram. Scale bar, 100 nm.

**Supplementary Figure 10.**
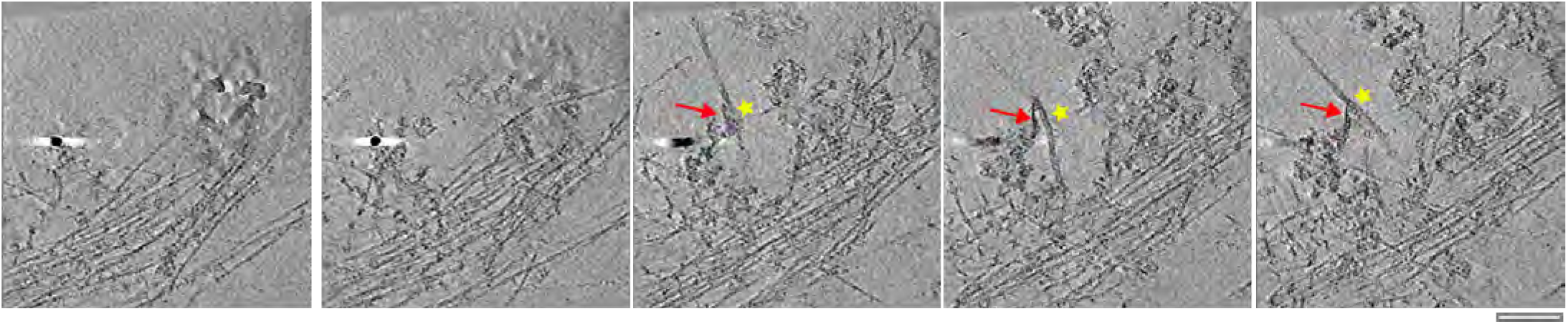
Tomographic slices showing a linear density bridging vimentin to FA-periphery densities. Sequential tomographic slices are shown from left to right, corresponding to increasing Z positions (from lower to higher along the Z axis). The linear density (red arrow) is observed linking vimentin filaments to FA-periphery densities across consecutive slices. The yellow star indicates the vimentin filament. Scale bar, 100 nm.

